# Chromatic topological mapping reveals organelle-specific spatial organization within microglia

**DOI:** 10.64898/2026.06.05.730307

**Authors:** Jens Agerberg, Lida Kanari, Margaret E. Maes, Wojciech Chachólski, Sandra Siegert, Martina Scolamiero

**Author notes:** Corresponding and last authors,.

## Abstract

In branched cells, including neurons and glia, intracellular organelles are distributed across complex cellular arbors where their spatial arrangement supports transport, signaling, and compartmentalized function. Although intracellular organelle organization is increasingly recognized as an important feature of cellular state and function, existing approaches assess organelle abundance or spatial position without accounting for the branching architecture that shapes cellular function. Here, we introduce the *chromatic topological morphology descriptor* (*chromatic TMD*), a framework that quantitatively resolves intracellular organization in relation to branching morphology. Applied to reconstructed microglia with annotated lysosomal and mitochondrial compartments across retinal layers and after optic nerve crush injury, *chromatic TMD* identifies distinct organelle-specific spatial programs: CD68^+^-endosomal-lysosomes undergo layer-dependent branch-specific redistribution, revealing selective intracellular reorganization after injury, whereas mitochondrial organization remains closely coupled to branching morphology. These findings establish intracellular organization as an additional layer of cellular architecture that can be systematically analyzed across branched neural cells.

## 1 Introduction

Neurons and glia exhibit highly branched morphologies that are central to how cells integrate into and interact within neuronal networks [1, 2]. Intracellular components, such as organelles, are distributed throughout these branched structures and support essential functions, including energy production, intracellular trafficking, and the degradation or recycling of cellular components [3, 4]. For example, in neurons, organelles must be transported between the soma and distal processes to ensure efficient distribution of energy and signaling information across long cellular distances [5]. Disruptions in this transport can impair neuronal activity and connectivity, contributing to neurological disorders [6]. Recent advances in microscopy now enable detailed visualization of the organelle network, together with its embedding within the entire branching tree [7, 8, 9]. Yet, quantification of organelles primarily focuses on their abundance, size, or distance from the cell soma [10]. This leaves several fundamental questions open about how intracellular organelles are arranged within branching cellular morphology, how this organization changes with functional state, and whether organelles actively reorganize along processes to support specific functions or simply scale with morphological remodeling. In microglia, a class of immune cells, these insights are critical, as they transition between surveillance and reactivity roles accompanied by pronounced remodeling of their branching morphology and their intracellular organization [11, 12]. For example, reactive microglia up-regulate the number of lysosomal organelles along cellular branches to support debris clearance and phagocytic activity [13, 14]. Similarly, mitochondrial organization along cellular branches can shift between extended, connected networks and isolated organelles, reflecting changes in cellular state and functional health [15, 16, 14]. Resolving cell-to-organelle relationships can reveal heterogeneity in intracellular organization among microglia and identify spatial signatures associated with different states, which conventional metrics do not preserve. To address this, we require methods that capture how organelles are organized within the cellular tree, rather than relying on measurements that treat morphology and intracellular organization separately.

Cellular architecture can naturally be represented as a tree-like structure embedded within a three-dimensional (3D) space. Topological data analysis (TDA) provides a principled mathematical framework to quantify such geometric structure. Specifically, the Topological Morphology Descriptor (*TMD*) algorithm provides a compact description of the tree [17], and has revealed differences in dendritic tree structure between pyramidal neurons in humans and mice [18], as well as enabled the separation of microglial morphologies across brain regions in healthy and pathological conditions [19, 14]. However, TMD focuses solely on tree morphology and lacks strategies to investigate the spatial organization of organelles within the tree. Chromatic topology, which studies spatial relationships among distinct entities [20], provides a natural mathematical framework but has not yet been specialized to the case of branching trees.

Here, we introduce *chromatic TMD*, which establishes relationships between the morphology tree and the organelle subgraph embedded within it. *Chromatic TMD* captures how organelles occupy, co-localize within, and are absent from branching cellular architecture while remaining computationally tractable. We apply this framework to reconstructed retinal microglia from the outer and inner plexiform layers (OPL and IPL) with annotated CD68^+^-endosomal-lysosomal and mitochondrial organelles under naïve conditions and following optic nerve crush (ONC), a well-established paradigm of retinal ganglion cell degeneration and microglial activation [21, 22, 23]. Because ONC induces pronounced CD68 upregulation in IPL microglia [14, 24, 25], this system enables comparison of how distinct organelle classes reorganize within branching microglial arbors across retinal layers and injury states. Together, our findings identify intracellular organization as an additional layer of microglial architecture shaped by organelle identity, retinal layer, and injury state.

## 2 Results

### 2.1 Microglia organelle distribution provides a model for *chromatic TMD*

Microglia are widely distributed across the brain, where they continuously survey and respond to local environmental changes. In the retina, microglia are spatially organized and selectively occupy the outer (OPL) and inner plexiform layers (IPL), where neuronal processes form synaptic contacts (**Figure 1A-C**). This well-defined laminar organization makes the retina a suitable system for studying layer-specific microglial responses to injury. Specifically, in the optic nerve crush (ONC) model, which induces reproducible perturbations in retinal ganglion cell axonal processes [21, 22, 23], microglial responses differ across layers in the ONC [14, 24, 25]. To investigate intracellular organization within this context, we used the Cx3cr1^CreERT2/+^×PhAM^fl/fl^ mouse model, which enables selective visualization of the mitochondrial network in microglia in combination with immunostaining of the endosomal-lysosomal marker CD68 and Iba1 labeling [14] (**Figure 1D**). This allows us to analyze CD68^+^-organelles and mitochondria within the same reconstructed microglial cells. We represent each microglia as a rooted tree, with vertices corresponding to spatial coordinates and edges representing the cell’s branching structure. In this representation, the root corresponds to the cell soma and the leaves to the terminal microglial process tips, marking the distal endpoints of the branching arbor. We define organelle-specific subgraphs by annotating vertices based on their overlap with reconstructed CD68 or mitochondria surfaces. This representation provides a structural framework for analyzing organelle organization relative to the cellular morphology.

**Figure 1:**
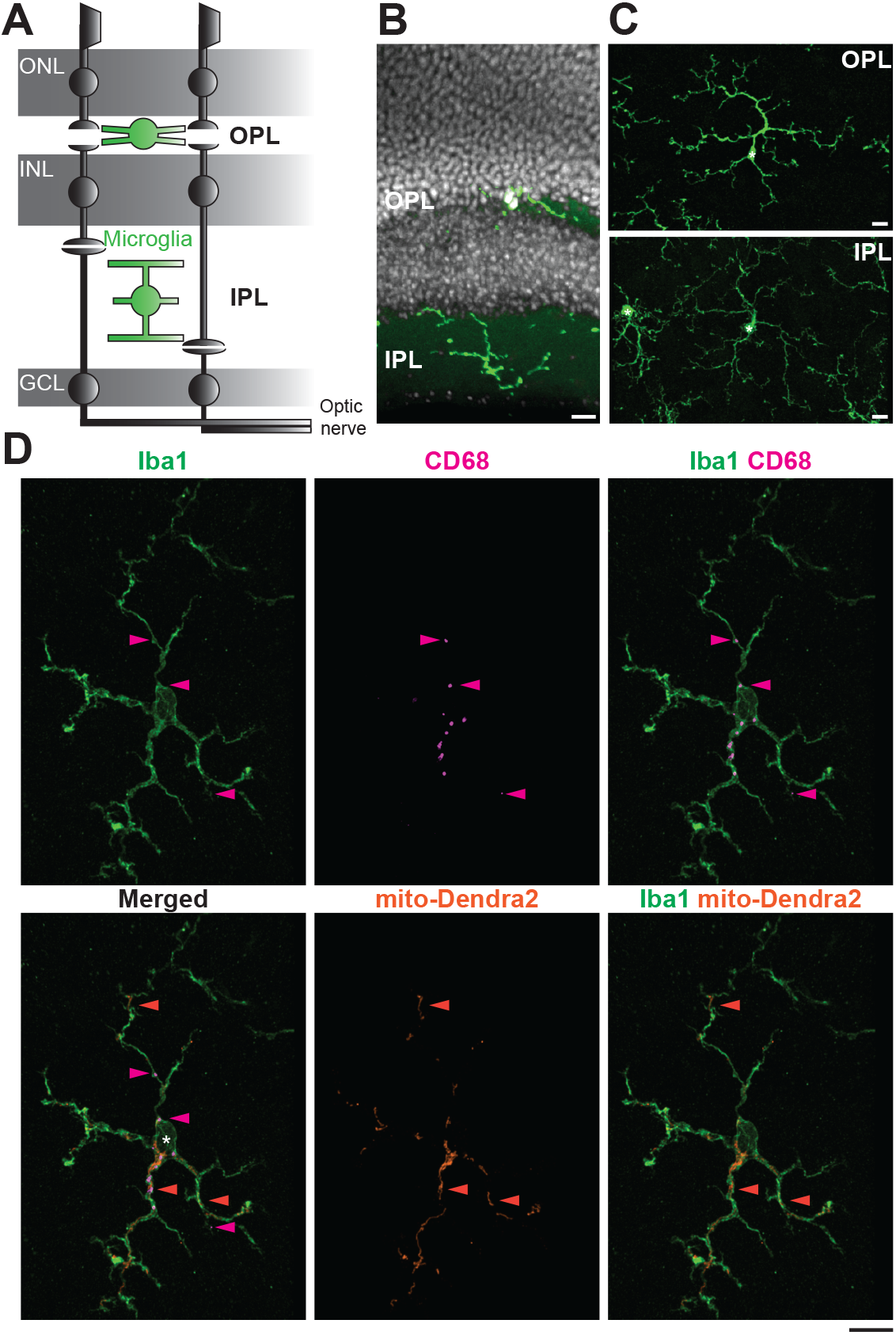
Organelle distribution in retinal microglia. **A**. Simplified side-view schematic of the retina emphasizing microglia (green) location in the synaptic outer plexiform layer (OPL) and inner plexiform layer (IPL). ONL, photoreceptor-containing outer nuclear layer. INL, inner nuclear layer. GCL, ganglion cell layer. Ganglion cell axons form the optic nerve. **B**-**D**. Immunostaining of microglia (Iba1, green). Scale bars: 10*µm*. **B**-**C**. Representative confocal images of microglia in OPL and IPL in retinal side-view with Hoechst dye-labeled nuclei (white, **B**), and whole mount view (**C**). **C-D**. Asterisks, soma location. **D**. Example microglia from adult, healthy Cx3cr1^CreERT2/+^×PhAM^fl/fl^ mouse model, with labeled organelles: CD68 as endosomal-lysosomal marker (magenta), PhAM/mito-Dendra2 for mitochondria (orange). Arrows: Positions of selected organelles within the microglial arbors.

### 2.2 Chromatic TMD captures organelle–tree relationships

Motivated by the application domain of microglia trees, we introduce the *chromatic TMD*, which establishes the relationship between the morphological tree *X* and the organelle subgraphs *Y* and how they are distributed across the tree’s branches (**Figure 2**). Specifically, we consider a rooted tree *X* with a path distance function *f* on its vertices and a subgraph *Y* ⊆ *X*, on which the path distance is defined by restriction. The function *f* defines super-level-set filtrations of the tree and subgraph, starting from the periphery and continuing until the root (soma) is reached. For a given *t* ∈ [0, ∞), the super-level set *X*_*t*_ = *f* ^−1^([*t*, ∞)) is the subgraph of *X* given by those vertices whose function values are greater than or equal to *t*, and edges between them. Super-level sets of the path distance define a filtration of *X* of the form 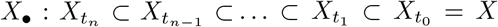, and a filtration of the subgraph 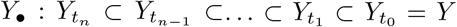 (**Figure 2B**). Next, we apply 0-th homology, with coefficients in a field, to the filtrations of *X* and *Y* . This determines a sequence of vector spaces and linear maps, denoted by *H*_0_(*X*_•_) and *H*_0_(*Y*_•_), which are also called *persistence modules*. Persistence modules admit a combinatorial description in the form of a barcode [26]. Since the distance-to-soma function *f* is order preserving with respect to the parent-child relation on the tree vertices, the barcode of *H*_0_(*X*_•_) equals the output of the *TMD* algorithm [27]. Similarly, the barcode of *H*_0_(*Y*_•_) describes the connected components of the organelle subgraph along the filtration. However, we cannot obtain the relation between *X* and *Y* by considering the filtrations of *X* and *Y* in isolation. Instead, we note that *Y*_*t*_ is a subgraph of *X*_*t*_, for every *t*, and that these inclusions determine an inclusion of filtrations *i* : *Y*_•_ ↪ *X*_•_ (**Figure 2B**). Our object of study is the map *H*_0_(*i*) : *H*_0_(*Y*_•_) → *H*_0_(*X*_•_) induced in 0-th homology by the inclusion of filtrations. Intuitively, at each step *t* of the filtration, this map encodes information about how the connected components of *Y*_*t*_ are related to the connected components of *X*_*t*_. In summary, *chromatic TMD* analyzes spatial relationships between the morphology tree and the organelle subgraph through the barcodes of five persistence modules: the tree morphology *H*_0_(*X*_•_), the organelle subgraph *H*_0_(*Y*_•_), the *image*, the *kernel*, and the *cokernel* of the map *H*_0_(*i*) (see Methods).

**Figure 2:**
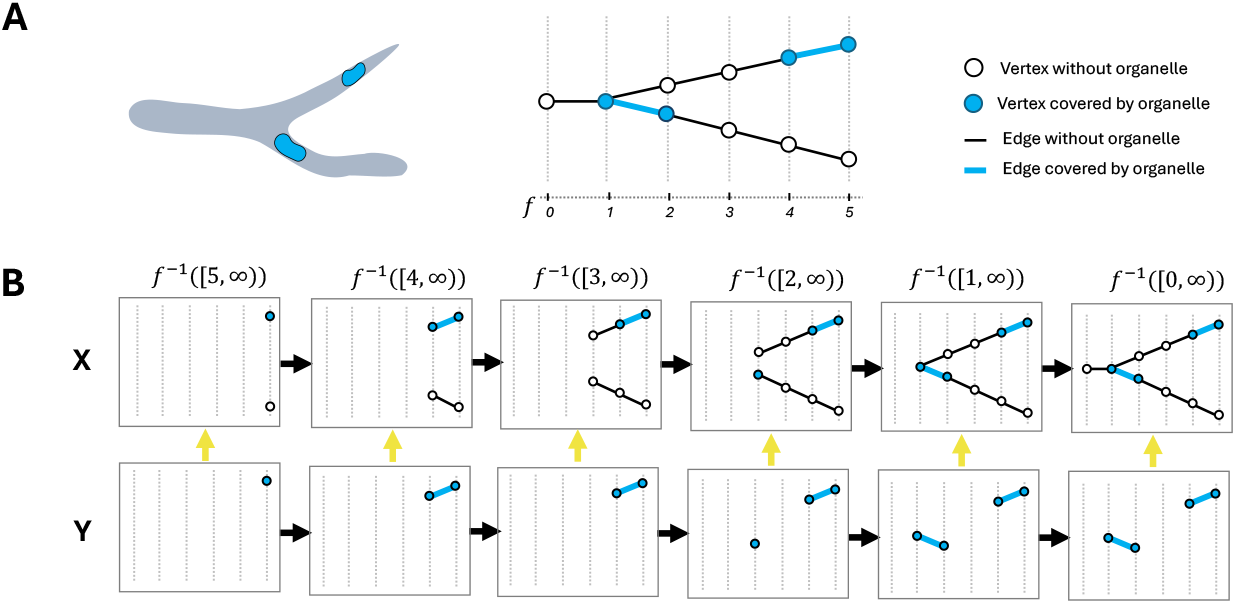
Filtration of a toy example tree with a subgraph. **A**. Left, schematic example of a branch (grey) containing organelles (blue). Next, skeletonization of the same branch consisting of edges and nodes along the path distance function *f* . **B**. Super-level set filtration based on path distance function *f*, leading to a point-wise inclusion of each vertex in its distance, resulting in super-level sets *Y*_*t*_ ⊂ *X*_*t*_. *X* (upper row) represents the tree, and *Y* (lower row) shows the organelles (blue vertices and edges) as a subgraph of the tree *X*. Yellow arrow: point-wise inclusion of super-level sets *Y*_*t*_ ⊂ *X*_*t*_.

### 2.3 Distributed and localized organelle–tree relationships with *chromatic TMD*

Organelle distribution is not uniform across a cell. They can be distributed across multiple branches or localized within a subset of branches (**Figure 1D**). To assess whether *chromatic TMD* captures these differences, we applied it to our toy example (**Figure 2A**), and organized the organelles either on separate (*distributed*) or on the same branch (*localized*, **Figure 3A**). The *TMD* barcodes of the branching tree are identical in both scenarios because the underlying morphology is unchanged (**Figure 3B**). Likewise, the persistence barcodes of the organelle subgraphs are identical because the organelles are positioned at the same distances from the soma in both examples. In contrast, the three invariants of *chromatic TMD* (**Figure 3B**) distinguish the two organizational patterns: When organelles are localized within the same branch, the *image* barcode collapses to a single bar. In contrast, the *kernel* barcode captures the co-localization of multiple organelles along the same branch, resulting in a longer bar in the localized case. Finally, the *cokernel* barcode identifies regions of the arbor lacking organelles and quantifies how far this absence extends along the branches. Together, these examples demonstrate that *chromatic TMD* resolves complementary aspects of how organelles occupy, co-localize within, and are absent from a branching tree.

**Figure 3:**
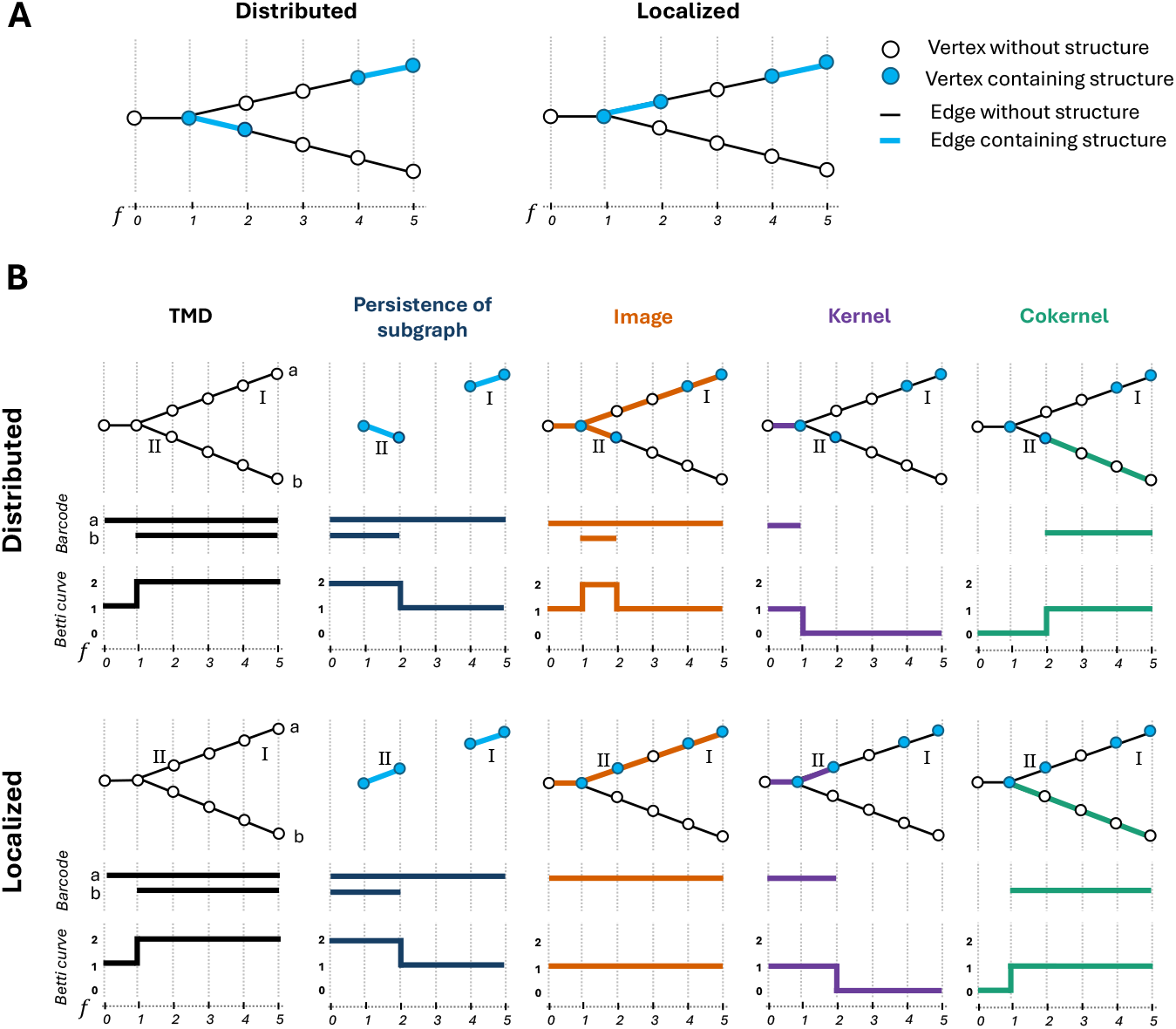
*Chromatic TMD* distinguishes organelle distribution in illustrative example. **A**. Skeletonization of the toy branch from **Figure 2A** consisting of edges and nodes along the path distance function *f* . The organelle distribution (blue) is shuffled on two different branches (distributed, left) or on the same branch (localized, right). **B**. Top row: distributed organelles. Bottom row: localized organelles. From left to right: TMD (topological morphological descriptor, black, *X*), persistence of subgraphs (blue, *Y*), and the three invariants of the *chromatic TMD* : *image* (orange), *kernel* (purple), and *cokernel* (green). Colors inside the skeleton highlight the different features for the two branches (a, b) and the subgraphs with organelles (I, II), followed below by the corresponding *barcode* and *Betti curve* representation. For *TMD*, each bar starts at a tip of the tree, with the branch either merging into another branch (bar [1, 5)) or being a primary process connecting to the soma (bar [0, 5)). For the *subgraph*, the bar end point corresponds to the path distance where the organelle first appears in the filtration, and the start point is either when two organelles connect by a branching point or is 0 if an organelle does not cover branching points of the tree (bars [0, 5) and [0, 2)). For the *image*, the bar endpoint corresponds to the path distance at which the organelle first appears, and the bar start either when the organelle merges into another one or when the branch containing the organelle merges into another branch with organelles in the tree (bar [1, 2) for the *distributed* scenario). For the *kernel*, the bar endpoint is the path distance at which the organelle appears on a branch where other organelles already exist, (bar [0, 2) for the *localized* scenario), or where two branches containing organelles merge (bar [0, 1) for the *distributed* scenario); the bar start point is always 0. For the *cokernel*, the bar end point corresponds to the path distance of the tip and the start either when the branch merges into another one (bar [1, 5) of the *localized* tree) or where an organelle appears on the branch (e.g., bar [2, 5) of the *distributed* tree).

Next, we applied *chromatic TMD* to the microglia tree containing CD68 organelles shown in **Figure 1D** (**Figure 4A**). First, the *TMD* barcode contains 28 bars, corresponding to the number of microglial terminal endpoints. Short bars reflect transient branches that rapidly merge into other branches, while longer bars reflect major branches. Bars that start at 0 indicate primary branches at the soma. The number of bars in the *subgraph* barcode starting at 0 corresponds to the total number of CD68 organelles on the tree (*n* = 12). The endpoints of these bars reflect the path distance at which the organelles are located. Short bars with non-zero start points indicate organelles located at branching points of the tree. The three invariants of the *chromatic TMD* further refine this distribution pattern: The *image* barcode detected 7 bars out of the 28 bars in the *TMD*, corresponding to branches containing at least one CD68 organelle. Each bar’s endpoint marks the path distance of the most distal organelle location. The first bar ends at 35.7, which is half of the maximum path distance of 73.7 based on the longest bar in the *TMD* barcode, suggesting that CD68 organelles localize closer to the soma than to distal branches. Based on the number of bars in the *subgraph* and the *image*, some branches must contain multiple organelles. The *kernel* barcode resolves this. The endpoint of the bar represents the path distance at which two branches containing organelles merge or at which an organelle appears on a branch where other organelles already exist.

**Figure 4:**
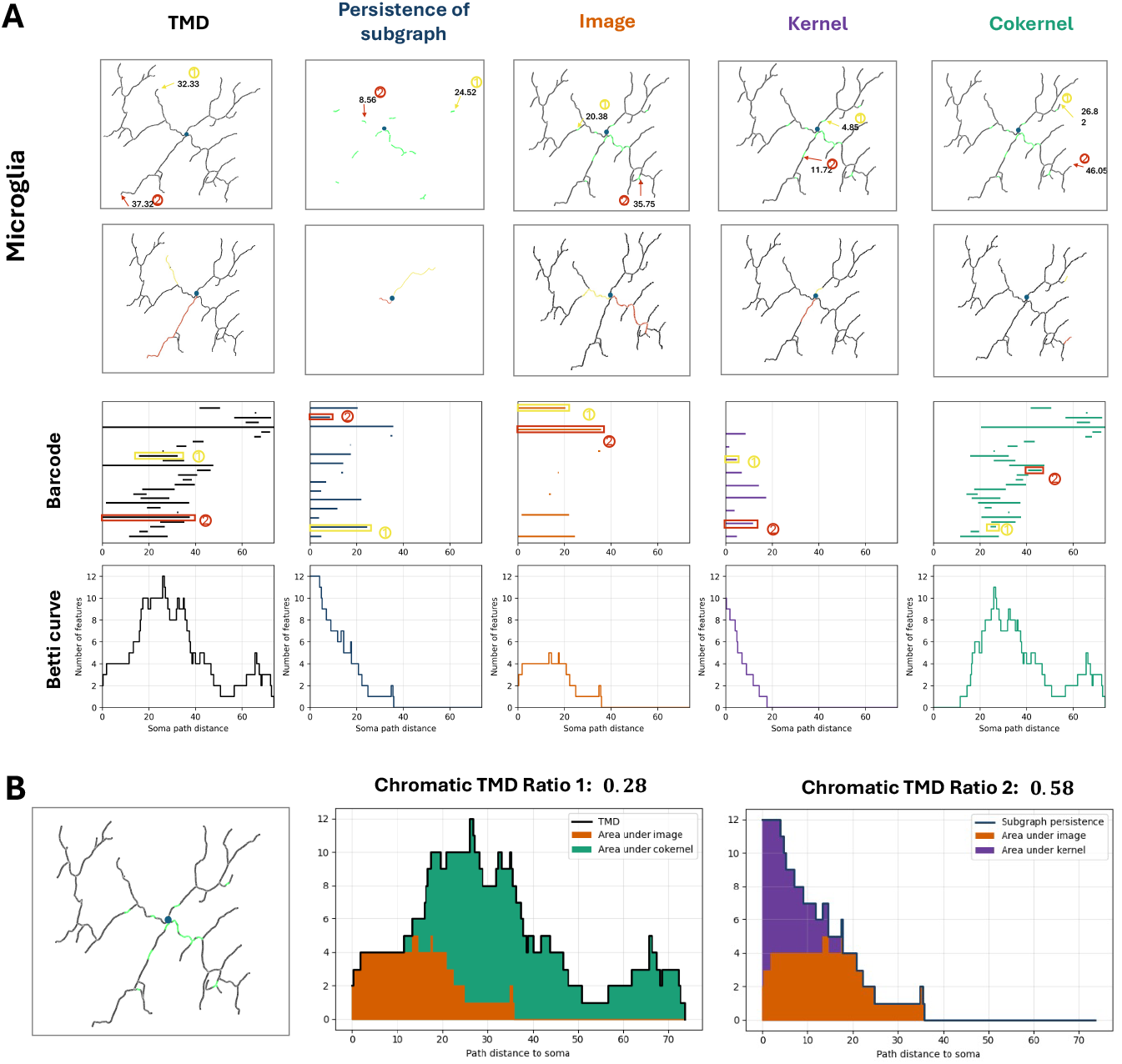
*Chromatic TMD* and ratios for the example microglia from Figure 1D. **A**. From left to right: TMD (topological morphological descriptor, black), persistence of subgraphs (blue), and the three invariants of the *chromatic TMD* : *image* (orange), recording the presence of organelles on branches, *kernel* (purple), capturing multiple organelle domains within individual branches, and *cokernel* (green), representing portions of the arbor that remain organelle-free. Top two rows: visual representations of a microglia branching tree (black) with soma (blue) and CD68 (green). Below, the barcode and Betti curves of the *chromatic TMD*. Each barcode highlights two example bars (yellow and red); their scale is represented in the tree representation corresponding to the bar endpoints (top row), and the path of the bar along the tree (below). **B**. The same microglia branching tree as in **A** (black) with labeled CD68 (green), and soma (blue). *Chromatic TMD ratio 1* derived as the decomposition of the TMD Betti curve into *image* (orange) and *cokernel* (green) and *ratio 2* derived as the decomposition of the subgraph persistence Betti curve into *image* and *kernel* (purple).

The presence of persistent bars in the *kernel* barcode suggests that CD68 organelles concentrate on relatively few branches. The bars in the *cokernel* barcode have the same number and endpoints as in the *TMD* barcode, while the bar length indicates how long the branch is absent from CD68. The similarity in bar lengths indicates that substantial portions of the arbor remain devoid of organelles. Combined with the organelle concentration on a few proximal branches, this pattern reflects a typical surveillance microglia.

### 2.4 Quantitative summaries of chromatic TMD barcodes

While barcodes provide detailed representations of how organelles are distributed along the branching structure of an individual cell, they are difficult to compare across populations. *Betti curves* summarize the structural information captured by the barcodes, reporting the number of bars present at each path distance (see Methods). For our microglia tree (**Figure 4A**), the *TMD* Betti curve corresponds to the number of branches present at path distance *t*. This is conceptually related to the Sholl analysis [28], but uses path distances rather than Euclidean distance. The Betti curve of the organelle *subgraph* reflects how many CD68 organelle components are present at path distance greater than or equal to *t*, showing the total number at scale 0 (**Figure 4A**). Extending this framework, the *chromatic TMD* Betti curves for *image, kernel*, and *cokernel* represent where CD68 organelles are located, how they concentrate on branches, and where CD68 is absent in branches, respectively. This framework enables further quantitative comparison and statistical analysis of how organelle-associated features relate to the overall branching tree. For this, we derived two *chromatic TMD* ratios from the *Betti* curves: an organelle coverage measure (*ratio 1*), quantifying how much of the arbor contains organelles, and an organelle distribution measure (*ratio 2*), capturing whether organelles concentrate on a few branches or distribute across the arbor (see Methods). For the microglia tree (**Figure 4B**), the low *ratio 1* emphasizes that the organelles occupy only parts of the tree, while their distribution across branches, as captured by *ratio 2* reflect a concentration of CD68 organelles on few branches. Together, these descriptors capture cellular morphology, organelle distribution, and their relationship, which we summarize in a cheat sheet for intuitive interpretation of typical patterns (**Supplementary Figure S1**).

### 2.5 Layer-specific CD68 redistribution accompanies microglial remodeling after ONC

To benchmark our *TMD* and *chromatic TMD* framework, we assessed microglia’s layer-specific response at baseline and upon retinal ganglion cell injury, using morphology and the spatial organization of CD68-positive organelles. First, we compared the microglia trees in the OPL and IPL under naïve, baseline conditions (**Figure 5A**). None of the commonly used morphometrics, such as the number of leaf nodes (terminal end points), the total process length, or the maximum Euclidean distance of a leaf to the soma, showed statistically significant differences between OPL and IPL (**Figure 5B**). In contrast, the *TMD Betti* curves indicated subtle adaptations, with a relative enrichment of proximal branches in the IPL and more distal branches in the OPL (**Figure 5C**). However, these differences were insufficient to separate the two populations at the single-cell level, as evidenced by the lack of clear clustering in the t-SNE embedding of the *TMD Betti* curves (**Figure 5D**). While the *TMD Betti* curve reports the number of bars at each distance, the *stable rank* curve summarizes the distribution of bar lengths in the TMD barcode (see Methods). Here, the largely overlapping mean *stable rank* curves indicate comparable branch-length distributions across layers (**Figure 5E**). When we examined the intracellular organization of CD68 organelles relative to the tree-branching structure, they were comparable across layers (**Figure 5F-G**). The *image* and *kernel* Betti curves showed no difference, while the *cokernel* had minor variation consistent with the morphological differences already reflected in the *TMD* Betti curves. This suggests that both the extent and distribution of CD68 occupancy along the arbor are largely indistinguishable between layers under baseline conditions, with organelles covering only a subset of branches (*ratio 1*) and remaining confined to a limited number of processes (*ratio 2*) (**Figure 5G**).

**Figure 5:**
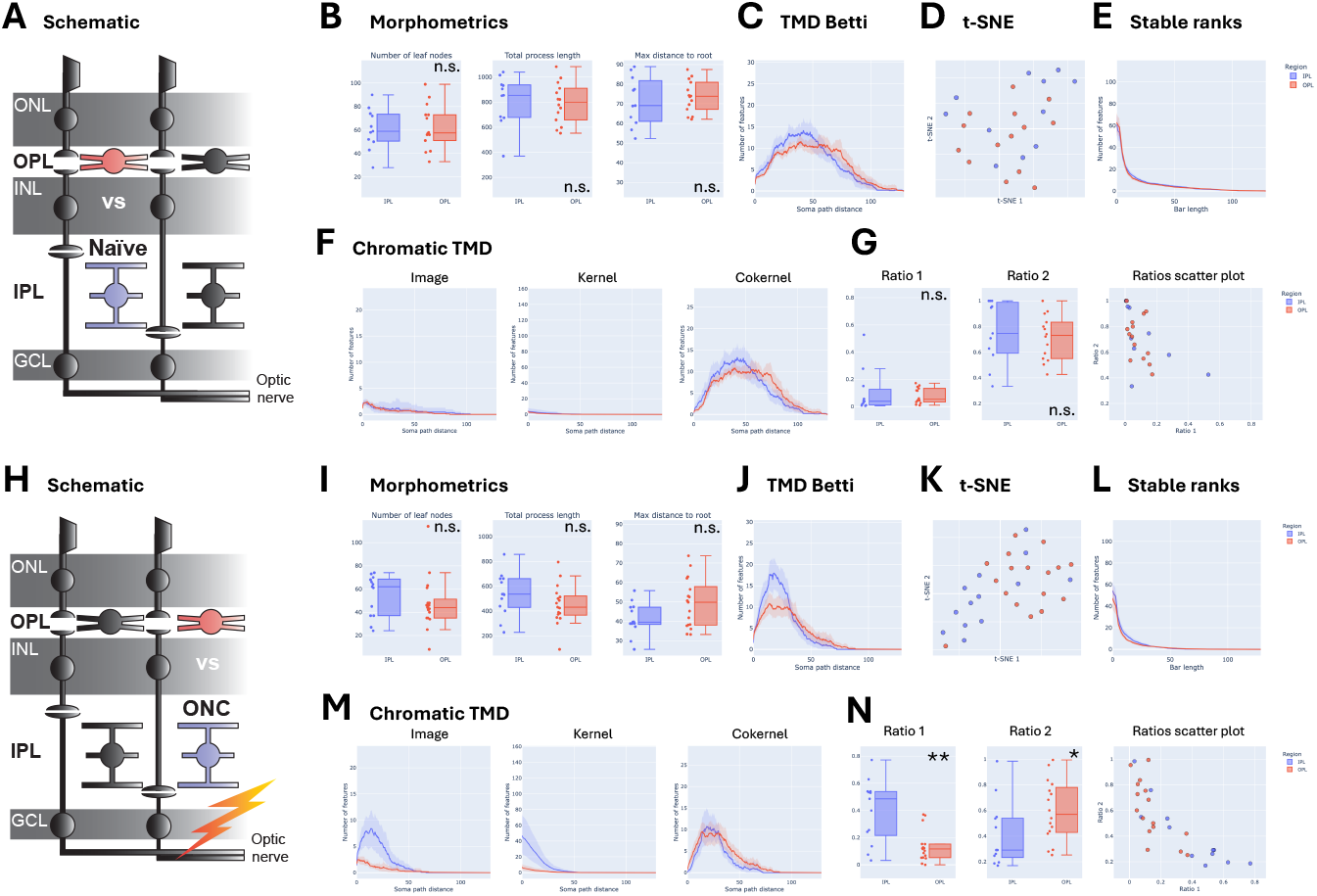
*Chromatic TMD* reveals layer-specific microglial reactivity and CD68 reorganization beyond classical morphometrics. Comparison between microglia in the outer (OPL, red, *n* = 25) and inner plexiform layer (IPL, blue, *n* = 30) for naïve (**A-G**) and after optic nerve crush (ONC, **H**-**N**). **A, H**. Schematic of the adult retina, side-view with the photoreceptor containing outer nuclear layer (ONL), inner nuclear layer (INL), and ganglion cell layer (GCL). Highlighted comparison. **B, I**. Box plots of *number of leaf nodes, total process length*, and *maximum distance of a leaf to root (soma)* with central line as median, inter-quartile range (Q1–Q3), and whiskers extending to the most extreme non-outlier values (1.5×IQR). Dots next to the box-plot, individual observations. Wilcoxon test for ONC *number of leaves*, all other: t-tests, ^ns^*p >* 0.05. **C, J**. *TMD* analysis with mean *Betti curves* showing number of features from soma path distance, with 95% confidence band. **D, K**. t-SNE plot. Each dot represents a single microglia, color-coded for OPL and IPL. **E, L**. Mean *stable ranks* curves representing number of bars with persistence longer than each threshold, with 95% confidence band. **F-G, M-N**. *Chromatic TMD* analysis of CD68. **F, M**. Mean Betti curves showing the number of features from soma path distance for *image, kernel*, and *cokernel* with 95% confidence band. **G, N**. Box plots of *ratio 1* and *ratio 2* with central line as median, interquartile range (Q1–Q3), and whiskers extending to the most extreme non-outlier values (1.5×IQR). Dots next to the box-plot, individual observations. t-test for naïve *ratio 2*, all other: Wilcoxon test, ^ns^*p >* 0.05, \**p <* 0.05, \*\**p <* 0.01. Next, scatter plot of *ratio 1 versus ratio 2*. Each dot represents a single microglia, color-coded by condition.

Next, we compared microglia in the OPL and IPL following optic nerve crush (ONC, **Figure 5H**). ONC induces microglial activation and increases CD68 levels within 5 days after injury, particularly in the IPL, where microglia are in contact with the apoptotic ganglion cells [14, 24, 25]. When we analyzed microglia from both layers using standard morphometric measurements, we found no difference (**Figure 5I**). In contrast, the *TMD* Betti curve of both layers peaked at approximately the same path distance, with the IPL having markedly higher *TMD* Betti values compared to the OPL, indicating an increased branching number proximal to the soma (**Figure 5J**). The t-SNE plot reflected partial separation of the cells by layer (**Figure 5K**), suggesting emerging differences in morphology at the single-cell level. The *stable rank* curves exhibited an increase in short-to medium-persistence features in the IPL, consistent with a shift toward shorter, more locally branched processes (**Figure 5L**). Together, these results indicate a layer-specific remodeling of microglial morphology in response to injury. The standard morphometric measures of CD68 abundance and size showed significant differences between OPL and IPL, whereas the distance from the cell soma did not (**Supplementary Figure S2**). The mean *image* and *kernel* Betti curves reflected a similar pronounced increase in the IPL compared with the OPL, but at the same time emphasized that more branches contained CD68 organelles and that they are more frequently organized within the same branch structures (**Figure 5M**). The *cokernel* Betti curves remained similar between layers, which in combination with the increased branching observed in the *TMD* Betti curve (**Figure 5J**), suggests that the additional formed proximal branches in the IPL are associated with increased CD68 organelles, while the extent of CD68-free branches remains largely unchanged between layers after ONC. This is further supported by a significant increase in organelle coverage of the arbor (*ratio 1*) (**Figure 5N**), whereas the lower *ratio 2* in IPL reflects that CD68 is more concentrated on the branches. This suggests increased organelle coverage on proximal branches, while some branches remain devoid of organelles, as indicated by the *cokernel*. Together, these results demonstrate that *TMD* and *chromatic TMD* resolve injury-induced, layer-specific CD68 reorganization within the microglial arbor, revealing branch-level changes not captured by common morphology parameters (**Figure 5B, I**).

### 2.6 Chromatic TMD identifies CD68 branch-level reorganization in IPL microglia

While the previous analysis established layer-specific differences, it does not resolve how microglia within each layer adapt to injury. To assess how microglia within each layer respond to the injury, we first compared microglial morphology between naïve and ONC conditions within each layer, starting with the IPL (**Figure 6A**). In the ONC condition, the mean *TMD* Betti curve shifted toward smaller path distances (**Figure 6B**). At the single-cell level, the t-SNE plot of the *TMD* Betti curves clustered in two well-separated groups corresponding to naïve and ONC cells (**Figure 6C**). Consistent with the *TMD* Betti curves, the *stable rank* curves preserved short-persistence branches but reduced medium- and long-persistence features under ONC (**Figure 6D**). Overall, the morphology reflects a reorganization toward shorter, more locally confined processes closer to the soma and reduced branch extension in the periphery following injury. In comparison, classical morphometrics yielded mixed results, with differences in the total process length and the maximum root-to-leaf distance, but not in the number of leaf nodes (**Supplementary Figure S3A**). This highlights the advantage of *TMD* in providing a consistent interpretation of structural changes across the arbor.

**Figure 6:**
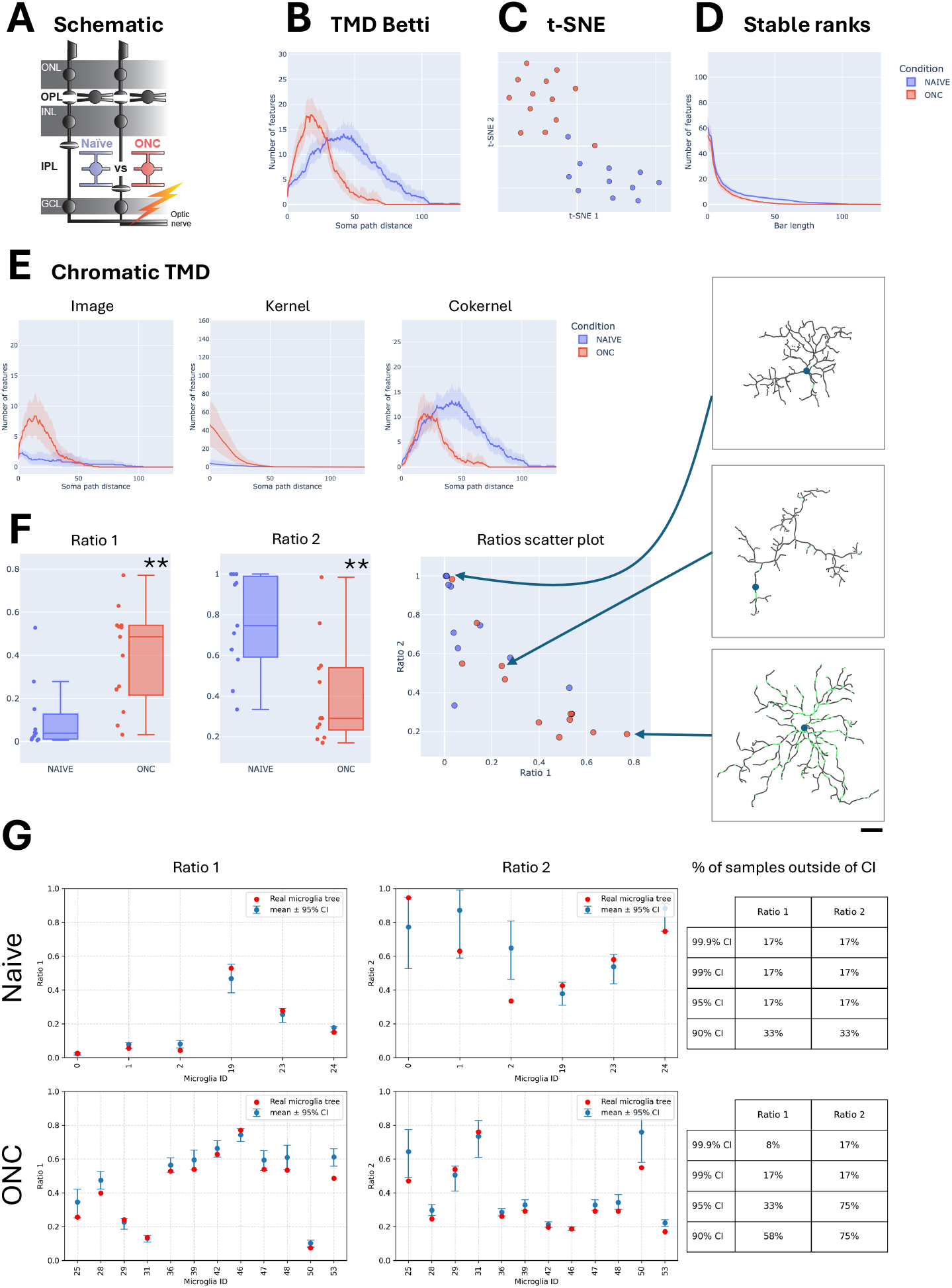
CD68 lysosome branch-level organization in microglia in IPL. Comparison of microglia in the inner plexiform layer (IPL) between naïve (blue, *n* = 11) and optic nerve crush (ONC, red, *n* = 13) conditions **A**. Schematic of the adult retina, side-view with the photoreceptor containing outer nuclear layer (ONL), inner nuclear layer (INL), and ganglion cell layer (GCL). Highlighted comparison. **B**. *TMD* analysis with mean*TMD* Betti curves showing number of features from soma path distance, with 95% confidence band. **C**. t-SNE of the *TMD* Betti curves. Each dot represents a single microglia, color-coded to the condition. **D**. Mean *stable ranks* curves representing the number of bars with persistence longer than each threshold, with 95% confidence band. **E-F**. *Chromatic TMD* analysis of CD68. **E**. Mean Betti curves showing the number of features from soma path distance for *image, kernel*, and *cokernel*, with 95% confidence band. **F**. Box plots of *ratio 1* and *ratio 2* with central line as median, inter-quartile range (Q1–Q3), and whiskers extending to the most extreme non-outlier values (1.5×IQR). Dots next to the box-plot, individual observations. Wilcoxon tests, \*\**p <* 0.01. Next, scatter plot of *ratio 1 versus ratio 2*. Each dot, a single microglia, color-coded to the condition with a highlighted microglia skeleton (black), soma (blue), and CD68 (green). Scale bar: 10*µm*. **G**. Null hypothesis of random assignments of CD68 organelles to individual microglia trees for *ratio 1* (left) and *ratio 2* (right) in naïve (top) and ONC (bottom) conditions. Red: observed *ratio* of the microglia ID with organelle distribution. Blue: mean and 95% confidence band for ratios *obtained* by randomizing the organelle distribution for the same microglial ID, with *n* = 1000. Next: Percentage of samples outside the confidence intervals.

Next, we focused on the distribution of CD68 as a function of morphological adjustment. Our analysis of classical morphometric parameters shows that an increase in CD68 abundance and size occurs, while the path distance remains unchanged under ONC compared to naïve (**Supplementary Figure S3 C-D**). Yet, the *chromatic TMD* for the ONC condition revealed that the maximum of the *image* Betti curve occurred at similar path distances as the *TMD* Betti curve (**Figure 6E**), indicating that despite a contraction of the tree morphology, CD68 organelles remain distributed at comparable distances to the soma. The marked increase in height relative to naïve, which showed only modest CD68 coverage, further emphasizes that several previously unoccupied branches become populated with CD68. The increase in the *kernel* Betti curve represents that multiple organelles are organized along the same branches. The shape of the *cokernel* Betti curve for ONC follows that of the naïve condition, indicating a similar extent of arbor regions devoid of CD68 close to the soma, whereas distally it vanishes under ONC, aligning with the contraction of the tree morphology. These changes in the *chromatic TMD* Betti curves are reflected in a significant increase in organelle coverage (*ratio 1*) after ONC, while the decrease in *ratio 2* supports a higher organelle occupation along the branches (**Figure 6F**). In the corresponding scatterplot, this joint behavior gives rise to a cluster of high *ratio 1* and low *ratio 2*, where the majority of ONC samples are found.

To further test whether this preferential concentration of organelles on a subset of branches arises from structural constraints of the underlying tree or does reflect specific patterns in CD68 organization, we compared the measured *ratio 1* and *ratio 2* to corresponding values on a null model in which CD68 organelles were randomly assigned to branches of the same microglial tree while preserving the distribution of their path distances to the soma (see Methods). The width of the resulting confidence intervals reflects the range of feasible organelle configurations under these constraints. For *ratio 1*, the confidence interval is narrow in both naïve and ONC conditions, indicating strong structural constraints on organelle placement (**Figure 6G**). In the naïve condition, the measured *ratio 1* values largely fall within the confidence interval; under ONC, most values shift below the interval. This suggests that organelles occupy fewer branches than expected under random placement. A similar pattern is observed for *ratio 2*, where 75% of the ONC cells fall below the 95% confidence interval, indicating that organelles are more strongly concentrated within a subset of branches than expected by chance. In contrast, in the naïve condition, only 17% of cells fall outside the interval, supporting an organelle distribution more consistent with a random placement, given the constraints of the underlying tree and the organelle path distances. Overall, these results demonstrate that ONC is associated with a branch-specific redistribution of CD68^+^-organelles within IPL microglia, resulting in a concentration of organelles relative to the morphology-preserving, branch-uniform null model.

### 2.7 OPL microglia undergo morphological remodeling with limited CD68 lysosomal reorganization

To assess whether microglia in the OPL undergo comparable injury-induced reorganization, we next examined morphological and CD68 changes in OPL microglia between naïve and ONC conditions (**Figure 7A**). Classical morphometric measurements showed significant reductions in the overall tree morphology (**Supplementary Figure S4B**), while CD68-related morphometrics remained largely unchanged (**Supplementary Figure S4D**). To resolve, beyond these global measures, how the branching structure and CD68 reorganize, we first applied *TMD*. The mean *TMD* Betti curve for ONC shifted toward smaller path distances but, in contrast to the IPL, did not increase in height (**Figure 7B**). This indicates that the overall branching complexity is largely preserved, while the arbor retracts toward the soma rather than undergoing structural expansion. At the single-cell level, the t-SNE plot of the individual *TMD* Betti curves similarly revealed two clusters corresponding to naïve and ONC conditions (**Figure 7C**). The mean *stable rank* curves under ONC showed fewer branches across all persistence scales, with the strongest reduction in short- and medium-persistence features (**Figure 7D**). To determine whether the retraction and shortening of the proximal branches in OPL microglia is accompanied by coordinated CD68 reorganization, we next analyzed the *chromatic TMD*. In contrast to the IPL, the *image* and *kernel* Betti curves revealed no significant ONC-induced changes (**Figure 7E**). The *cokernel* Betti curve shifted toward smaller path distances while maintaining a similar height, mirroring the shift observed in the *TMD* Betti curve (**Figure 7B**). Both *ratios* remained unchanged between naïve and ONC microglia. This suggests that despite substantial morphological remodeling, the relative spatial organization of CD68 lysosomes remained largely preserved within the branching tree. Finally, we tested whether the effects arise from structural constraints of the underlying tree or reflect specific patterns in CD68 organization. For *ratio 1*, the confidence interval remained narrow in both naïve and ONC conditions, indicating that feasible organelle coverage is constrained by the morphology and the small organelle number (**Figure 7G**). In the naïve condition, all measured values fell within the 95% confidence interval, consistent with largely random CD68 branch occupancy. Following ONC, however, 29% of the cells were outside the confidence interval, with most values trending toward below the expected range. This suggests that a subset of OPL microglia exhibits reduced CD68 branch occupancy following injury, compared with a random distribution of organelles on the branches, although this effect is less pronounced than in the IPL. For *ratio 2*, confidence intervals were broader overall, indicating greater flexibility in how organelles can be distributed across branches. In naïve OPL microglia, only 10% of cells deviate from the null model. After ONC, however, 36% of cells fell outside the confidence interval, with most values similarly biased toward the lower half of the expected range. This indicates that ONC promotes partial clustering of CD68 organelles within fewer branches in a subset of cells, but without the coordinated branch-specific lysosomal redistribution observed in the IPL. Overall, these findings demonstrate that OPL microglia undergo pronounced morphological remodeling after injury, but CD68 organization remains comparatively stable. This contrasts with the coordinated branch-specific lysosomal reorganization observed in the IPL, supporting a layer-specific microglial response to ONC.

**Figure 7:**
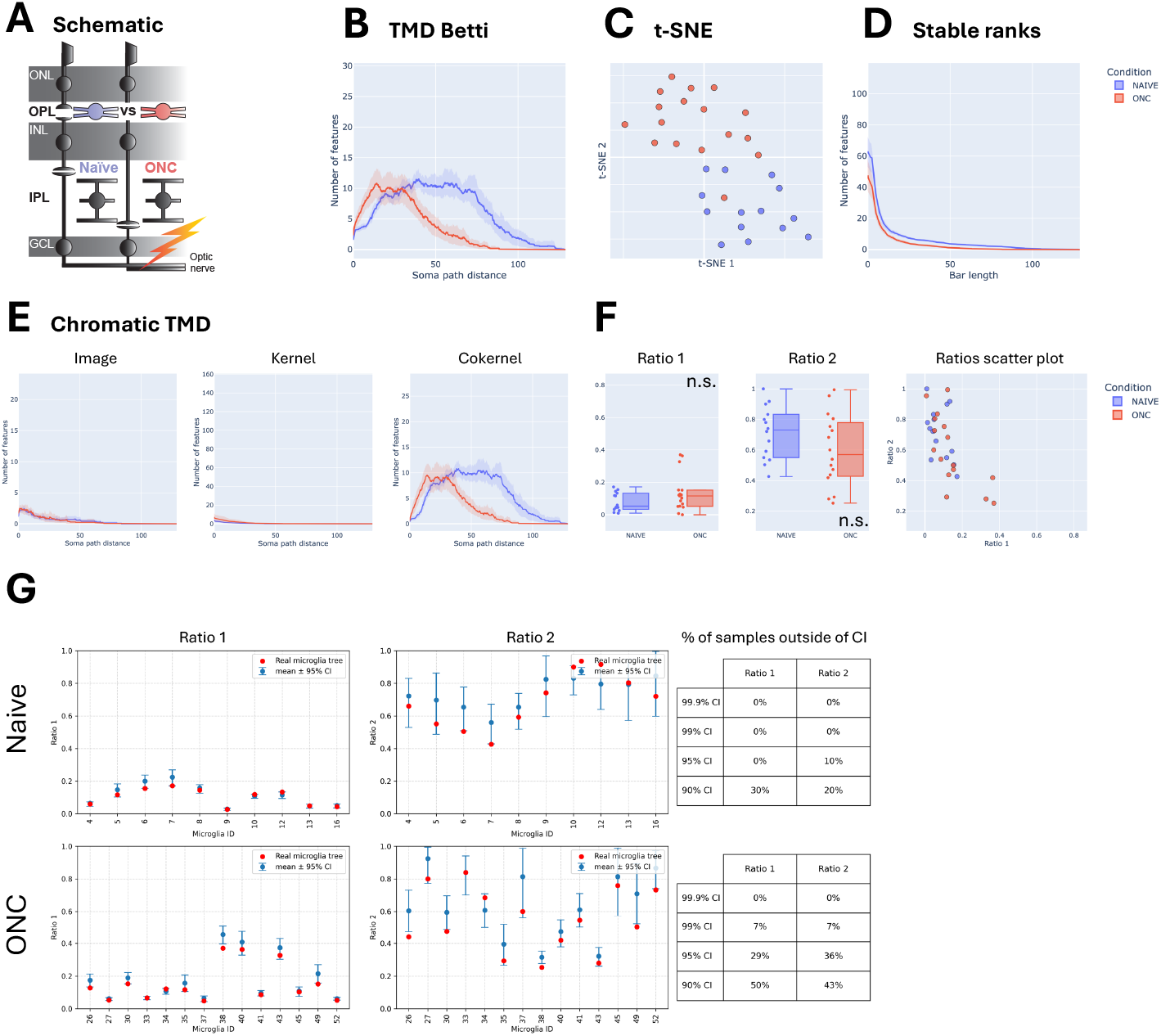
OPL microglia morphological adaptation with limited effects on CD68 constraints. Comparison of microglia in the outer plexiform layer (OPL) between naïve (blue, *n* = 14) and optic nerve crush (ONC, red, *n* = 17) conditions. **A**. Schematic of the adult retina, side-view with the photoreceptor containing outer nuclear layer (ONL), inner nuclear layer (INL), and ganglion cell layer (GCL). **B**. Morphological description of the microglia trees with *TMD Betti* curves (**B**) t-SNE of the *Betti curves* (**C**), and *stable ranks* (**D**). **B**. *TMD* analysis with mean *TMD Betti curves* showing number of features from soma path distance, with 95% confidence band. **C**. t-SNE plot of the *TMD* Betti curves. Each dot represents a single microglia, color-coded to the condition. **D**. Mean *stable ranks* curves representing the number of bars with persistence longer than each threshold, with 95% confidence band. **E-F**. *Chromatic TMD* analysis of CD68. **E**. Mean Betti curves showing number of features from soma path distance for *image, kernel*, and *cokernel* with 95% confidence band. **F**. Box plots of *ratio 1* and *ratio 2* with central line as median, inter-quartile range (Q1–Q3), and whiskers extending to the most extreme non-outlier values (1.5×IQR). Dots next to the box-plot, individual observations. Wilcoxon test for *ratio 1*, t-test for *ratio 2*, ^ns^*p >* 0.05. Next, scatter plot of *ratio 1 versus ratio 2*. Each dot represents a single microglia, color-coded by condition. **G**. Null hypothesis of random assignments of CD68 organelles to individual microglia trees for *ratio 1* (left) and *ratio 2* (right) in naïve (top) and ONC (bottom) conditions. Red: observed *ratio* of the microglia ID with organelle distribution. Blue: mean and 95% confidence band for ratios *obtained* by randomizing the organelle distribution on the same microglia ID for *n* = 1000. Next: Percentage of samples outside the confidence intervals.

### 2.8 Mitochondria organization exhibits stable organelle-morphology coupling across layers and injury conditions

Having established that *chromatic TMD* resolves layer- and injury-specific lysosomal reorganization, we next asked whether this framework similarly captures the spatial organization of mitochondria, a distinct organelle system with complementary functional roles in microglial physiology. Classical mitochondrial morphometrics detected injury-associated changes in abundance, size, and spatial extent (**Supplementary Figure S5A**). To capture the coupling between mitochondrial organization and branching architecture, we applied *chromatic TMD* to mitochondrial distributions within the same reconstructed microglial cells, enabling direct comparison under identical morphological conditions (**Figure 8A**). Across both retinal layers, mitochondrial *chromatic TMD* profiles revealed a broadly similar ONC-associated organizational pattern. In both layers, the mitochondrial mean *image, kernel*, and *cokernel* Betti curves shifted toward smaller path distances following ONC, indicating that mitochondrial organization largely follows the proximal arbor remodeling induced by injury (**Figure 8B-C**), In the OPL, mitochondrial *image* occupancy remained largely stable, while the *kernel* component decreased, suggesting that mitochondrial components become redistributed along more proximal branches without major changes in overall branch occupancy (**Figure 8B**). Similarly, in the IPL, mitochondrial organization also shifted proximally, with only modest changes in branch occupancy (**Figure 8C**). Thus, within the same microglial morphologies that showed pronounced CD68 reorganization, mitochondrial organization instead scaled in proportion to injury-induced structural remodeling. When we quantified the *chromatic ratios*, we found that in the OPL, *ratio 1* remained unchanged while *ratio 2* increased significantly after ONC (**Figure 8 D**). This indicates that mitochondrial branch coverage remains stable, but mitochondrial domains become more selectively organized across the arbors. Similarly, in the IPL, *ratio 1* showed minimal change, whereas *ratio 2* also increased significantly after ONC (**Figure 8E**). This consistent increase in *ratio 2* across both layers suggests that, within the same injury-remodeled cells, mitochondrial organization becomes more strongly aligned with branch architecture without substantial expansion of total branch coverage. To determine whether these mitochondrial patterns arise from structural constraints of the underlying tree, we compared observed ratios with null models preserving mitochondrial number and path-distance distribution (**Figure 8F-G**). In contrast to CD68, confidence intervals for both *ratio 1* and *ratio 2* were consistently narrow across all layers and conditions, and observed values systematically fell below null-model expectations. In OPL microglia, 93% of the naïve cells and 71% of ONC cells were outside of the 95% confidence band for *ratio 1*, while 100% of naïve and 94% of ONC cells deviated for *ratio 2* (**Figure 8F**). We observed similar proportions in the IPL (**Figure 8G**). These demonstrate that mitochondrial placement deviates from the branch-uniform null model and follows a characteristic pattern, even after controlling for the underlying microglial morphology and mitochondrial distance to the soma. Although ONC modestly reduced the proportion of cells outside the null model, mitochondria organization remained strongly non-random across all conditions. Together, these findings demonstrate that mitochondrial organization remains strongly coupled to branching topology in microglia across layers and injury conditions, in contrast to the dynamic, layer-dependent reorganization observed for CD68^+^-organelles.

**Figure 8:**
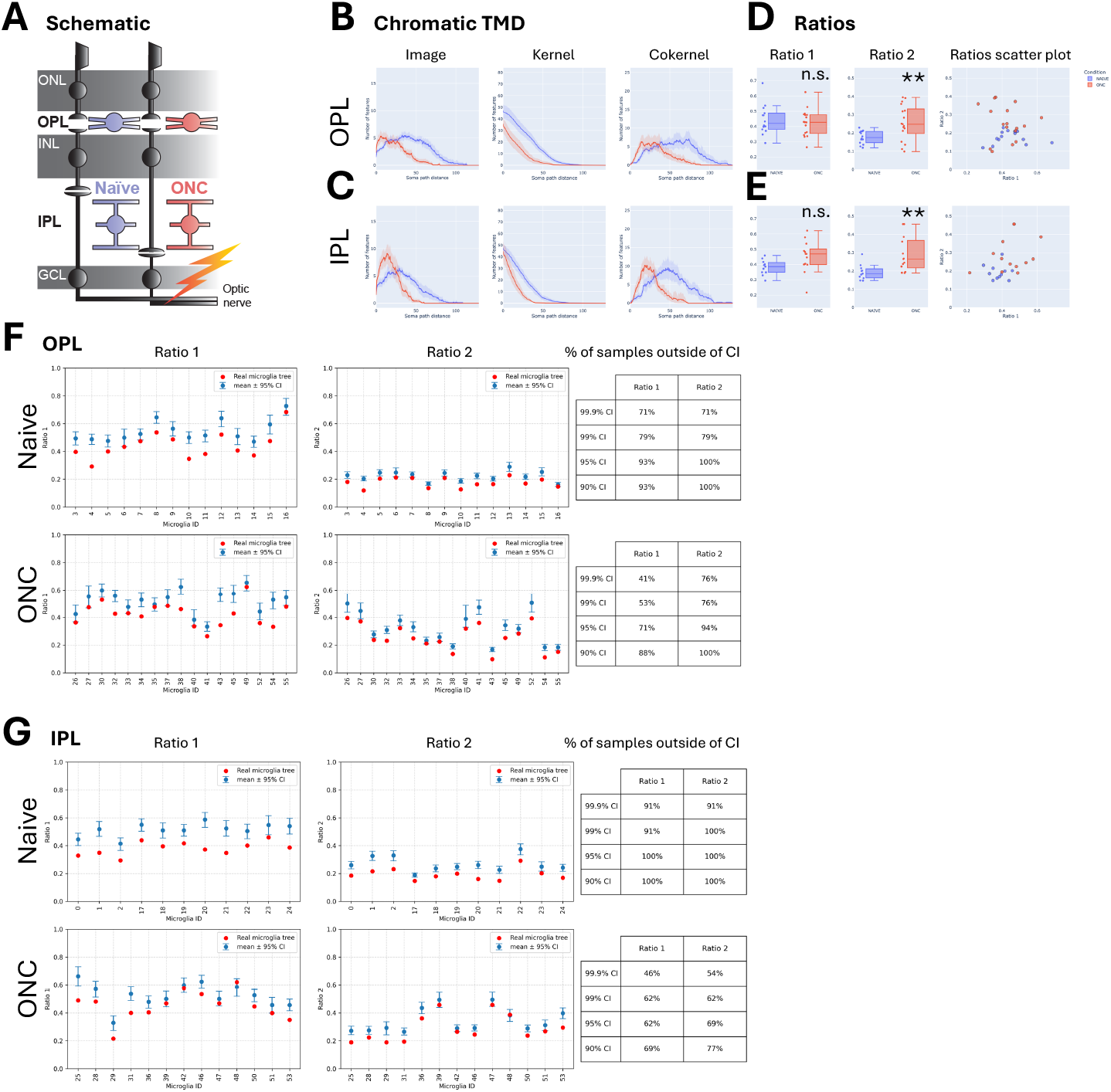
Mitochondria distribution is constrained by the microglia morphology. **A**. Schematic of the adult retina, side-view with the photoreceptor containing outer nuclear layer (ONL), inner nuclear layer (INL), and ganglion cell layer (GCL). Comparison between microglia in the OPL and IPL (outer- and inner plexiform layer, respectively) in naïve (blue, *n* = 27) *versus* optic nerve crush (ONC, red, *n* = 24) condition. **B-E**. *Chromatic TMD* analysis of mitochondria in OPL (**B, D**) and IPL (**C, E**). **B**-**C**. Mean *Betti curves* showing number of features from soma path distance for *image, kernel*, and *cokernel* with 95% confidence band. **D**-**E**. Box plots of *ratio 1* and *ratio 2* with central line as median, inter-quartile range (Q1–Q3), and whiskers extending to the most extreme non-outlier values (1.5×IQR). Dots next to the box-plot, individual observations. t-tests, ** *p <* 0.01. Next, scatter plot of *ratio 1 versus ratio 2*. Each dot represents a single microglia, color-coded by condition. **F**-**G**. Observed values, mean and confidence bands under assignment of organelles to random branches of *ratio 1* (left), and *ratio 2* (right) in naïve (top) and ONC (bottom) conditions for single microglia. Red: *ratio* of the microglia with actual organelle distribution. Blue: mean and 95% confidence band for ratios obtained by randomizing the organelle distribution on the microglia. Next: Percentage of samples outside the confidence intervals.

## 3 Discussion

The spatial organization of intracellular compartments is an important but difficult-to-quantify dimension of cellular architecture in branching cells. Here, we introduce *chromatic TMD* as a topology-based framework that directly links intracellular organization to cellular morphology. By resolving the distribution of organelles relative to branching morphology, *chromatic TMD* establishes intracellular organization as an additional dimension of cellular architecture. Combining mathematical rigor with an experimentally accessible interpretative framework (**Supplementary Figure S1**), *chromatic TMD* positions intracellular topology as a quantifiable organizational principle.

Reactive microglial remodeling has classically been described as a process of retraction and arbor simplification. Our findings refine this view by demonstrating that injury-induced remodeling involves a systematic reorganization of the branching topology rather than merely global contraction. *TMD* Betti curves and *stable rank* analyses showed that optic nerve injury redistributes microglial branching architecture toward shorter, more proximal, and less persistent branches (**Figure 6, 7**). Importantly, this remodeling differed by retinal layer: IPL microglia increased proximal branching complexity, whereas OPL microglia primarily exhibited arbor retraction. These results already demonstrate that topological descriptors capture systematic structural remodeling beyond what conventional morphometrics can capture. This has important implications for intracellular organization, which *chromatic TMD* can now capture. We show that organelles engage in distinct spatial programs during injury: In IPL microglia, ONC induced coordinated branch-specific redistribution of CD68^+^-organelles, characterized by increased branch occupancy, enhanced within-branch clustering, and strong deviations from random expectation (**Figure 6**). In contrast, OPL microglia displayed pronounced morphological remodeling but weaker and more heterogeneous lysosomal redistribution (**Figure 7**). This is consistent with localized intracellular reorganization in the local tissue context, which is not captured in simple abundance measurements.

A major biological and conceptual advance of this study is the demonstration that distinct intracellular systems can follow fundamentally different spatial programs within the same cellular morphology. Because we analyzed mitochondria and CD68^+^-organelles within identical reconstructed microglial cells, *chromatic TMD* enabled direct organelle-specific comparison while controlling for shared morphology. While lysosomes underwent dynamic, state-dependent redistribution, mitochondrial organization remains strongly coupled to branching topology across layers and injury conditions, closely mirroring overall arbor remodeling rather than independent organelle-specific redistribution (**Figure 8**). These findings are consistent with prior studies showing that microglial mitochondria occupy structurally constrained positions [14, 16] and support a model in which mitochondria preserve constitutive, architectural organization, whereas lysosomes exhibit dynamic state-dependent spatial organization. Taken together, these findings suggest that intracellular organization follows organelle-specific structural programs rather than a single shared organizational principle. This distinction, now possible with *chromatic TMD*, is particularly important because conventional abundance-based measures cannot determine whether organelles are broadly deployed throughout the arbor or selectively concentrated within specific branches. Two cells may contain similar organelle abundance yet differ profoundly in how those organelles are structurally deployed, potentially reflecting distinct functional states. *Chromatic TMD* directly resolves this limitation by distinguishing proportional scaling from genuine branch-level spatial reorganization.

For TDA, *chromatic TMD* is a new, versatile, and computationally efficient tool. Classical persistent homology and *TMD* consider only global morphology [29, 27], and recent chromatic TDA approaches primarily focus on relationships between cell populations as colored point clouds. Such chromatic topological approaches include geometric constructions such as Dowker complexes [30], Witness complexes [31], and Discrete Chromatic Structures [32], as well as topological summaries within chromatic TDA [20]. However, these methods do not explicitly model how intracellular structures are embedded within branching morphologies. *Chromatic TMD* instead directly tackles this problem, providing a linear-time algorithm compared to cubic-complexity algorithms in the standard case. The integration of *image, kernel*, and *cokernel* descriptors with interpretable *ratios* and null-model testing provides both rigorous mathematical structure and practical biological interpretability that remain inaccessible to traditional analyses. More broadly, these findings align with growing evidence across branching biological systems that organelle positioning follows non-random structural principles. Recent neuronal connectomic studies similarly showed that mitochondrial localization is constrained by neurite architecture and local structural features [10]. *Chromatic TMD* complements such approaches by providing a global topology-based framework that captures branch occupancy, internal compartmentalization, and organelle-free domains across the full cellular architecture. Thus, this framework broadens intracellular spatial biology from local positional analyses toward system-level structural organization.

Several limitations should be considered when interpreting these findings. First, in the ONC paradigm, retinal layer identity and distance to the lesion site are correlated, such that the observed differences between IPL and OPL microglia cannot fully dissociate injury-associated effects from intrinsic layer-specific microglial biology. Second, organelle reconstruction depends on imaging resolution and segmentation fidelity, particularly for smaller or densely packed organelles. Although persistent homology is theoretically stable under small perturbations, we did not systematically evaluate the robustness of *chromatic TMD* to segmentation uncertainty. Future studies using higher-resolution modalities such as expansion microscopy or dynamic live-cell imaging may further refine these analyses. In addition, the current framework analyzes one organelle system at a time. Extending *chromatic TMD* to jointly model multiple intracellular organelle systems and their relationships will be advantageous for identifying the interplay between different organelles. Third, the current morphology-preserving null model also does not incorporate additional structural constraints such as branch caliber or branch-order weighting, which may further refine expected occupancy distributions. Finally, broader application across additional cell types and pathological contexts will be necessary to determine how general these principles are across spatial cell biology.

Together, our study establishes *chromatic TMD* as both a mathematical framework and a biologically interpretable tool for quantifying intracellular organization within branching cells. By revealing intracellular topology as a previously inaccessible layer of cellular organization, *chromatic TMD* provides a foundation for topology-aware spatial cell biology and offers broad utility for studying how subcellular architecture contributes to structural specialization, adaptive remodeling, and cellular state across diverse biological systems.

## 4 Methods

### 4.1 Microglia Data Collection

#### 4.1.1 Animal procedures

Microglia reconstructions were sampled from preparations of a previous study in the Siegert laboratory (see: [14]). In short: *Cx3cr1* ^CreERT2^ (#020940) [33] were crossed to PhAM^fl/fl^ (#018385) [34]. Freshly prepared tamoxifen (Sigma Aldrich, T5648-5G) was administered via *intraperitoneal* injection daily (150mg/kg) for three days. Optic nerve crush (ONC) experiments were performed at adult animals at least three weeks after tamoxifen induction when monocyte populations have turned over [35]. Retinas were collected 5 days after ONC. All animal procedures are approved by the Bundesministerium für Wissenschaft, Forschung und Wirtschaft (bmwfw) Tierversuchsgesetz 2012, BGBI. I Nr. 114/2012, idF BGBI. I Nr. 31/2018 under the numbers 2021-0.607.460, 2023-0.250.844, 2025-0.781.719, 2020-0.272.234, and BMWFW-66.018/0005-WF/V/3b/2016.

#### 4.1.2 Immunostaining

Staining was carried out as described in [14]. In short, retinas were always protected from light to preserve the endogenous mito-Dendra2 fluorophore. After blocking the tissue, retinas were exposed to the following primary antibodies: rabbit anti-Iba1 (GeneTex, Cat#GTX100042, Lot 41556, RRID: AB 1240434 1:750) and rat anti-CD68 (Bio-Rad, Cat#: Cat#: MCA1957, RRID: AB 322219, 1:500). After incubation for three nights at 4°C on an orbital shaker, retinas were exposed to secondary antibodies for 2 hours, and to the nuclei dye Hoechst 334422 (1:5,000, Thermo Fisher Scientific, H3570) for 10 min, followed by washing and mounting.

#### 4.1.3 Confocal microscopy

Single cell images for deconvolution were acquired on Zeiss LSM800 microscope using a Plan-Apochromat 40× objective NA 1.4 (#420762-9900) at 1.5× digital zoom to acquire a pixel size of 0.05 µm in X and Y resolution and 0.150 µm in Z. Images were acquired from the peripheral region of the retina, in two opposing quadrants, and inner plexiform layer microglia were selected by setting the z-stack between the ganglion cell and inner nuclear layer base don the Hoechst staining. After IPL selection, the Iba1 channel was visualized, and the nearest Iba1^+^-cell was selected for imaging, thereby reducing bias in cell selection and blinding to the cell’s mitochondrial architecture. Tile scans (2×2) were necessary to acquire images of single IPL microglia under naïve conditions, resulting in images with dimensions of 202×202 µm. For each animal, 2-3 images from two opposing quadrants were acquired. Images were stitched using the Zen 2.3 or 3.5 desktop version.

#### 4.1.4 Image processing

SVI Huygens Professional v21.10 (https://svi.nl/HomePage) was used for batch deconvolution processing of stitched confocal images. Microscope parameters for batch settings were as follows: 50 nm sampling intervals for X and Y, 150 nm in Z. The refractive indices of the embedding medium were set to 1.515 and 1.49, respectively. Objective quality was listed as ‘good.’ For channel settings, the back-projected pinhole was set to 250 nm, and the excitation fill factor was 2. Excitation (ex) and emission (em) settings for the four channels were as follows: 405/420 nm, 488/520 nm, 561/602 nm, 633/650 nm. Deconvolution parameters were set to the default settings for the ‘confocal low template’ using the classic MLE (maximum likelihood estimation) algorithm with a maximum of 30 iterations. Images were output in the IMARIS file format. All confocal images presented in the figures were masked in IMARIS using the Iba1-immunostained microglia of interest to remove microglia and mitochondrial labeling that did not belong to the analyzed cell. Supplementary videos were animated using IMARIS.

#### 4.1.5 Mitochondria and CD68 surfaces in single microglia

Surface rendering analysis was performed using IMARIS v9.1-9.3 (Bitplane, IMARIS). Individual cells were cropped from the original tiled image when necessary to reduce processing time. Then, 3-dimensional surfaces were created for microglia, mitochondria, and CD68 vesicles using the *Surface* wizard with surface details settings of 0.5 µm, 0.2 µm, and 0.4 µm, respectively. Each surface was manually edited to resolve any inconsistencies in surface creation. Mitochondrial surfaces were carefully edited to verify connectivity throughout the network, often requiring manual splitting of surfaces that were joined by the thresholding within the wizard.

#### 4.1.6 Mitochondrial organelle and CD68 vesicle localization in single microglia

Completed *Surfaces* for either mitochondria or CD68 were ‘split’ using the Xtension *Split Surfaces* available via Bitplane IMARIS (https://imaris.oxinst.com/open/). This generates a *Split Surface* group, in which each individual organelle/vesicle becomes a new, uniquely named *surface* nested within the *Split Surface* group.

A *spot* 5 µm in diameter was manually added to identify the nucleus. Then, the *Spots and Surfaces distance* XTension calculates the distance between the nucleus *spot* and the center point of each individual *surface* in the *Split surface* group. The Xtension was modified from the original version to allow for up to 100 distance calculations and to output a .*csv* file.

#### 4.1.7 Reconstruction of microglia

After filtering and background subtraction, images were imported into IMARIS 9.2.v (Bitplane IMARIS). Microglial processes were traced with the filament-tracing plugin. Since the filament-tracing plugin provides a semi-automated reconstruction, this eliminates the need for a user-blind approach for selecting representative microglia. New starting points were detected when the largest diameter was set to 12 µm, and the seeding point size to 1 µm. Disconnected segments were removed with a filtering smoothness of 0.6 µm. After the tracing, we manually removed cells that were sitting at the border of the image and were only partially traced, so that these cells would not be analyzed. The generated skeleton images were converted from .ims format (IMARIS) to .swc format 157 by first obtaining the positions (x,y,z) and the diameter of each traced microglial process using the ImarisReader toolbox for MATLAB (https://github.com/PeterBeemiller/ImarisReader) and then exporting for format standardization using the NL Morphology Converter (http://neuroland.org). Artifacts from the 3D reconstructions failed to convert into SWC format automatically. A total number of 55 microglia cells were collected from either age-matched naïve or ONC animals from either the OPL or the IPL, including also cells already used in [14]. The core structure of the skeleton was encoded as a rooted tree with coordinates in physical space being recorded for each vertex. Vertices were annotated to indicate if they reside in regions of high concentration of either mitochondria or CD68, forming two subgraphs of the rooted tree (see 4.2.2). The data describing the tree structure of microglia, vertex positions, and labels for CD68 and mitochondria are stored in an extended SWC format, generated from the IMARIS filaments with an IMARIS Python Extension (https://git.ista.ac.at/iof-group/image-analysis/iof-pyimaris). The output SWC file contains an additional column with label IDs linking to IMARIS Surfaces. Features of IMARIS Surfaces are exported in separate tab-separated files. Path distance of all vertices to the root vertex is computed with the Python Navis library.

### 4.2 Persistent homology, TMD and summaries

#### 4.2.1 Persistent homology

Persistent homology studies the evolution of homology along a filtration of simplicial complexes *{K*_*t*_*}*_*t*∈ℝ_. For each homological dimension *k*, the inclusion maps *K*_*s*_ ↪ *K*_*t*_ for *s* ≤ *t* induce linear maps *H*_*k*_(*K*_*s*_) → *H*_*k*_(*K*_*t*_), yielding a sequence of vector spaces and linear maps known as a *persistence module*. Under suitable finiteness conditions, such modules admit a decomposition into interval modules, giving rise to a barcode representation [26]. In this representation, each interval encodes the birth and death of a *k*-dimensional topological feature. These barcodes provide a stable, multiscale summary of the intrinsic topological structure of the data.

#### 4.2.2 Tree function pairs

Let *X* = (*V*_*X*_, *E*_*X*_) be a rooted tree, a connected acyclic graph with a distinguished vertex called the root. The rooted tree structure defines a parent-child relation on *V*_*X*_ where any vertex in the tree, except the root, has exactly one parent. The parent of *v* ∈ *V*_*X*_ is the vertex *w* such that (*v, w*) ∈ *E*_*X*_ and *w* is on the unique path from *v* to the root. If *w* is parent to *v*, we say that *v* is a child of *w* and denote it by *w < v*. Indeed the parent-child relation provides the vertex set of *X* with a poset structure (*V*_*X*_, *<*) with the root being the minimal element. Let *f* : (*V*_*X*_, *<*) → ℝ be a an order preserving function, for *u, v* ∈ *V*_*X*_, *u* being a parent of *v* implies *f* (*u*) *< f* (*v*). We call (*X, f*) a *tree function pair*.

##### Example 4.1

*An order preserving function f* : *V*_*X*_ → ℝ *can be chosen to be the path distance to the root in X. If X is a tree embedded in an Euclidean space and f* (*v*) *is defined as the Euclidean distance of the vertex v to the root in X, the function is instead not necessarily order preserving*.

The function *f* : *V*_*X*_ → ℝ can be extended to edges in the graph, by defining the values on edges to be the minimum value of the vertices that it connects, that is *f* ((*u, v*)) = min(*f* (*u*), *f* (*v*)) for (*u, v*) ∈ *E*_*X*_ . Through this extension we have defined a map *f* : *X* → ℝ where *X* is interpreted as a simplicial complex. Moreover *f* has the property that for all edges (*u, v*), *f* ((*u, v*)) ≤ *f* (*u*) and *f* ((*u, v*)) ≤ *f* (*v*). This implies that the superlevel sets of the function *f*, {*f* ^−1^[*t*, ∞)} _*t*∈ℝ_ satisfy {*f* ^−1^[*t*, ∞)} ⊆ {*f* ^−1^[*s*, ∞)} when *t* ≥ *s* and form a filtration of spaces.

#### 4.2.3 Topological Morphology Descriptor (TMD)

The TMD algorithm [17] is applied to a tree function pair (*X, f*). The output is a barcode: a multiset of intervals on the real line. In the case when *f* is order preserving, it corresponds to the barcode decomposition of the 0-th persistent homology (see Section 4.2.1) of the superlevel set filtration of the function *f* on the tree *X* [27]. A formulation of the TMD as a recursive tree algorithm (useful for comparing it to the *Chromatic TMD*) is provided in Algorithm 1. The algorithm is initialized by being called on the root vertex.

#### 4.2.4 Betti curves

Betti curves are functional summaries of a barcode. If a barcode is represented by a Betti curve *f*, then *f* (*t*) corresponds to the number of bars where the start point is smaller or equal than *t* and the end point is larger than *t*. If the TMD filtration function is the Euclidean distance to the root then the Betti curve of the TMD corresponds to the Sholl descriptor [28], which tracks the number of branches that intersect a circle of radius *t* centered on the soma [27].

#### 4.2.5 Stable ranks

A stable rank [36] is a functional summary of a barcode in the form of a non-increasing piecewise constant function with values in [0, ∞). After choosing a pseudometric between barcodes, the stable rank of a barcode *X* corresponds to tracing the minimal rank (number of bars) of barcodes in increasing neighborhoods of *X* with respect to the chosen pseudometric. With the standard choice of pseudometric (corresponding to the bottleneck distance [37]), if a barcode is represented by a stable rank *f*, then *f* (*t*) corresponds to the number of bars with length greater or equal than *t*.

### 4.3 Chromatic TMD

#### 4.3.1 Filtrations of pairs of spaces

We consider a rooted tree *X* = (*V*_*X*_, *E*_*X*_) and a subgraph *Y* = (*V*_*Y*_, *E*_*Y*_) of *X*, i.e. *V*_*Y*_ ⊂ *V*_*X*_, *E*_*Y*_ ⊂ *E*_*X*_ and *u, v* ∈ *V*_*Y*_ if (*u, v*) ∈ *E*_*Y*_ . If *X* is a labeled tree, *Y* may be obtained by taking all vertices and edges with a particular label. Note that while *Y* is a subgraph of *X* it is not necessarily a connected graph.

Consider now superlevel sets of the function *f* : *X* → ℝ and of the restriction *f* |_*Y*_ of *f* to *Y* . We let *s* = (*s*_1_, *s*_2_ … *s*_*m*_), where *{s*_1_, *s*_2_, …, *s*_*m*_*}* = *{f* (*v*) : *v* ∈ *V*_*X*_ *}*

##### Algorithm 1 TMD

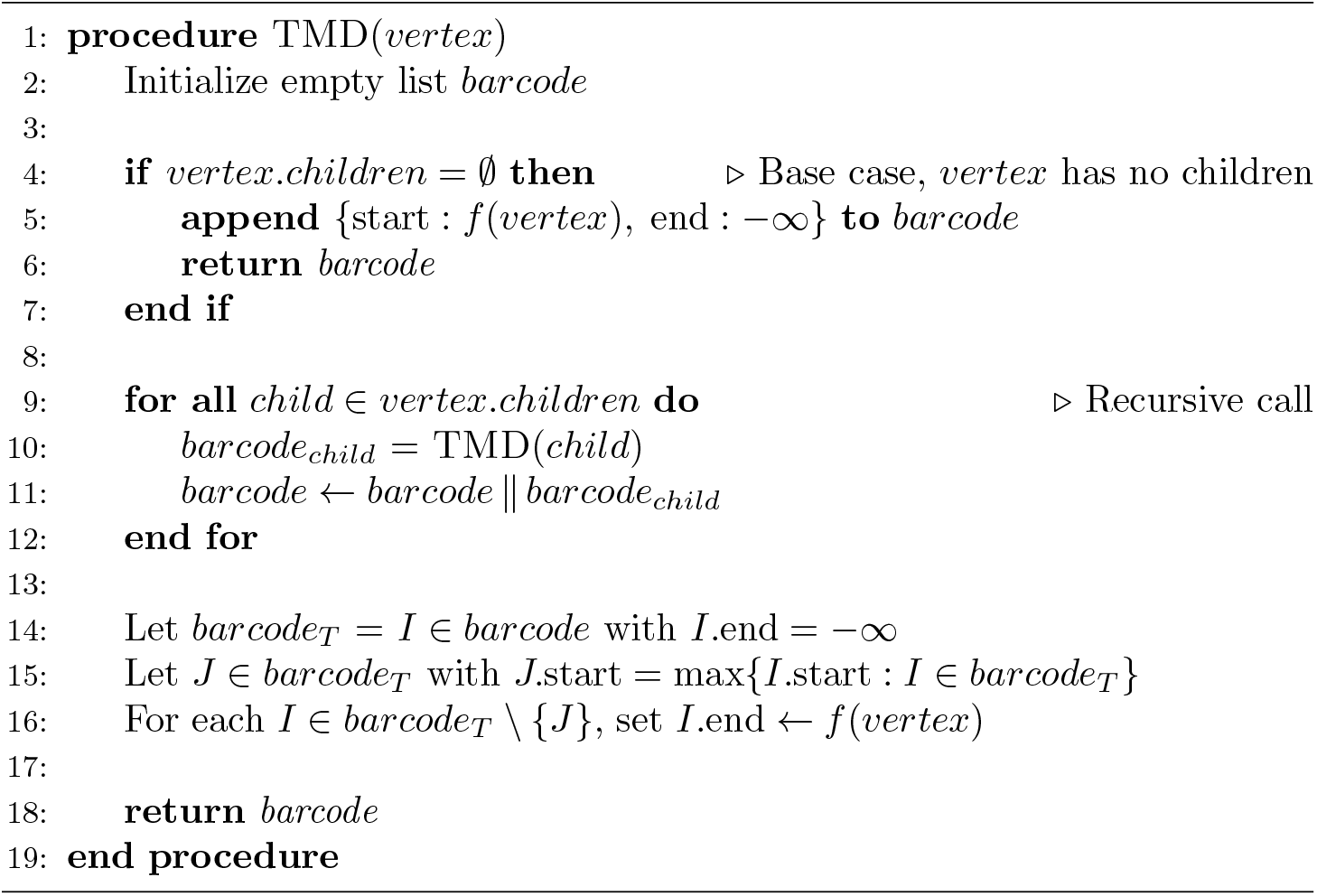

and *s*_1_ *< s*_2_ *<* · · · *< s*_*n*_. We let *X*_*i*_ = *f* ^−1^([*s*_*i*_, ∞)) and 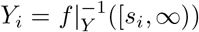. For every *i* we have *X*_*i*_ ⊆ *X*_*i*−1_, *Y*_*i*_ ⊆ *Y*_*i*−1_ and *Y*_*i*_ ⊆ *X*_*i*_. We can thus consider the commutative diagram of 0-th homology groups and maps induced by inclusions:

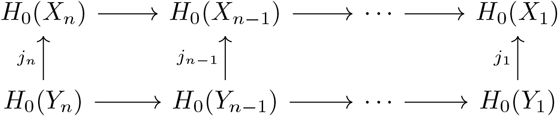

We are in a special case of the setting described in [38]: the *image, kernel* and *cokernel* of the map {*j*_*i*_} _*i*_, with *j*_*i*_ : *H*_0_(*Y*_*i*_) → *H*_0_(*X*_*i*_) are persistence modules and can be decomposed into barcodes. Contrary to [38] however, the decomposition can be computed with a recursive tree algorithm (Algorithm 2) in the spirit of the TMD (Algorithm 1).

#### 4.3.2 Algorithm

Given a tree-function pair (*X, f*) and a subgraph *Y* of *X*, the *chromatic TMD* is a linear-time complexity recursive tree algorithm to compute the barcode decomposition of the *image, kernel*, and *cokernel* persistence modules of the map *H*_0_(*i*) : *H*_0_(*Y*_•_) → *H*_0_(*X*_•_). The *chromatic TMD* can be seen as a generalization of the TMD and recovers this algorithm when the organelle subgraph coincides with the microglia tree (*image H*_0_(*i*) = *H*_0_(*X*_•_)) and when no organelle is present (*cokernel H*_0_(*i*) = *H*_0_(*X*_•_)). To describe the recursive structure, we call *X*(*w*) the subtree of *X* rooted at a vertex *w*, and *Y* (*w*) = *X*(*w*) ∩ *Y*, which is the restriction of the organelle subgraph to *X*(*w*). When we consider the superlevel set of *X*(*w*) by *f*, for a value slightly larger than *s* = *f* (*w*), we obtain a space that is homotopic to the union of all trees rooted at the children of *w*. That is, if *v*_1_, …, *v*_*n*_ are the children of *w*, then there exists *t > s* such that 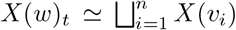, while *X*(*w*)_*s*_ = *X*(*w*). Since *X*(*v*_*i*_), *X*(*v*_*j*_) are disjoint spaces for all *i, j* = 1, …, *n* and *i* ≠ *j*, we have 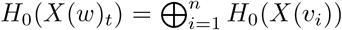 and 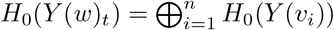. The persistence barcodes for *image, kernel*, and *cokernel* of the spaces *X*(*w*), *Y* (*w*) can thus be fully described in terms of the barcodes *image, kernel*, and *cokernel* of *X*(*v*_*i*_), *Y* (*v*_*i*_) for *i* = 1, …, *n* together with the transition *t* ≤ *s*. This allows the *chromatic TMD* algorithm to be formulated as a recursive tree algorithm, described by the base case when *w* is a leaf node, without children, and by the transition *t* ≤ *s*. The algorithm is initialized by being called on the root vertex. The pseudocode is provided in Algorithm 2, and a proof of correctness is given in Supplementary Section B.

### 4.4 Quantification and statistical analysis

#### 4.4.1 Data compilation and metric calculations

Python version 3.11 with packages *numpy, pandas, plotly, matplotlib* were used for data compilation, visualization and analysis. For dimensionality reduction, *L*_2_ distances between TMD Betti curves were computed and embedded into 2D with the t-SNE dimensionality reduction algorithm [39] using *scikit-learn*.

#### 4.4.2 Morphometrics

All morphometrics are computed based on the extended SWC files: The *number of leaf nodes* of the tree (tips of the microglia) corresponds to the count of terminal vertices, i.e. vertices with no children. The *total process length* is the sum of the lengths of all edges in the tree, providing a measure of overall arbor extent. The *maximum distance to the root* is the largest Euclidean distance between the root (soma) and any vertex, capturing the maximal spatial extent of the microglia. The *number of components* is the number of connected components in the subgraph, reflecting how fragmented or clustered the distribution of CD68 or mitochondria is along the tree structure. The *components process length* is the total length of all edges within these components, quantifying the overall extent of marker-associated processes. The *components process length (%)* is the components process length divided by total process length. The *components path distance* is the average of the distances of the marked vertices to the root along the tree, capturing how the marker is distributed with respect to the underlying arbor.

#### 4.4.3 CD68 and mitochondria occupancy bar chart

For each microglia tree, the processes of the tree are segmented and assigned to bins based on their path distance to the soma, using intervals of 0 − 10%, …, 90 − 100% where 100% corresponds to the maximum path distance of any vertex in the tree. For each bin, the fraction of process length covered by CD68 or mitochondria is computed. The bar chart reports the mean of these fractions across all microglia within each condition.

#### 4.4.4 Chromatic TMD ratios

From the *Betti* curves corresponding to the *image, kernel* and *cokernel* of the *chromatic TMD*, two ratios are derived: an organelle coverage measure (*ratio 1*, and an organelle distribution measure (*ratio 2*). *Ratio 1* is defined as the integral of the *image* Betti curve divided by the integral of the *TMD* Betti curve and *ratio 2*, defined as the integral of the *image* Betti curve divided by the integral of the Betti curve of the subgraph *Y* (**Supplementary Figure S6A-B**). These measurements relate to organelle occupancy based on the linear equations

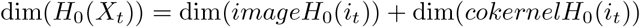

and

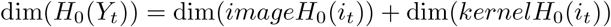

implying that the sum of the *image* and the *cokernel Betti* curves equals the *TMD Betti* curve and that the sum of the *image* and the *kernel* Betti curves equals the subgraph *Y* Betti curve.

#### 4.4.5 Null model based on synthetic placement of organelles

For each microglial tree, the connected components (organelles) of the subgraph corresponding to mitochondria or CD68 are extracted. For each component, we record (i) the path distance of the vertex closest to the soma and (ii) the process length covered by the component. The synthetic placement algorithm uses this information as input. For each component, a branch is selected uniformly at random among those that do not already contain a component at the corresponding path distance, and the component is assigned to that branch. Components are processed in random order until all have been placed. This yields a new subgraph of the microglial tree with the same number of components, preserving both their path distances and process lengths, but with components redistributed across branches. By repeating this procedure many times and computing *ratio 1* and *ratio 2* for each realization, we obtain a distribution of these ratios that defines a null model for random component placement on the tree. The ratios of the observed data can then be compared to this distribution.

#### 4.4.6 Statistical analysis

Statistical tests were performed using the Python package scipy. Groups for comparison were first tested for normal distribution and equal variance using the Shapiro test and Levene test, respectively. In the normal case Student t-test was used and otherwise Wilcoxon rank-sum test was used. Significance levels are indicated using the following notation (^ns^*p >* 0.05; \**p <* 0.05; \*\**p <* 0.01; \*\*\**p <* 0.001). Data are shown at the single-cell level, with cells sampled from at least three animals per condition. To assess whether group-level patterns were dominated by individual animals, we also inspected animal-level summaries, which showed consistent directionality across animals. A detailed statistical analysis can be found in the **Supplementary Table 1**.

## Acknowledgments

We thank the scientific service units at ISTA, the imaging optic facility (IOF) and specific Christoph Sommer for writing the extended .swc format, the pre-clinical facility (PCF), specifically Sonja Haslinger; the Siegert team members for constant feedback on the project.

## Author contributions

Conceptualization, Methodology: JA, LK, MEM, WC, MS, SS; Software: JA; Validation, Investigation: JA, MS, SS; Formal Analysis: JA; Writing – Original Draft: JA, MS, SS; Visualization: JA, SS; Supervision: WC, MS, SS; Funding acquisition: SS and MS. All authors reviewed and approved the final manuscript.

## Supplementary figures

**Figure S1:**
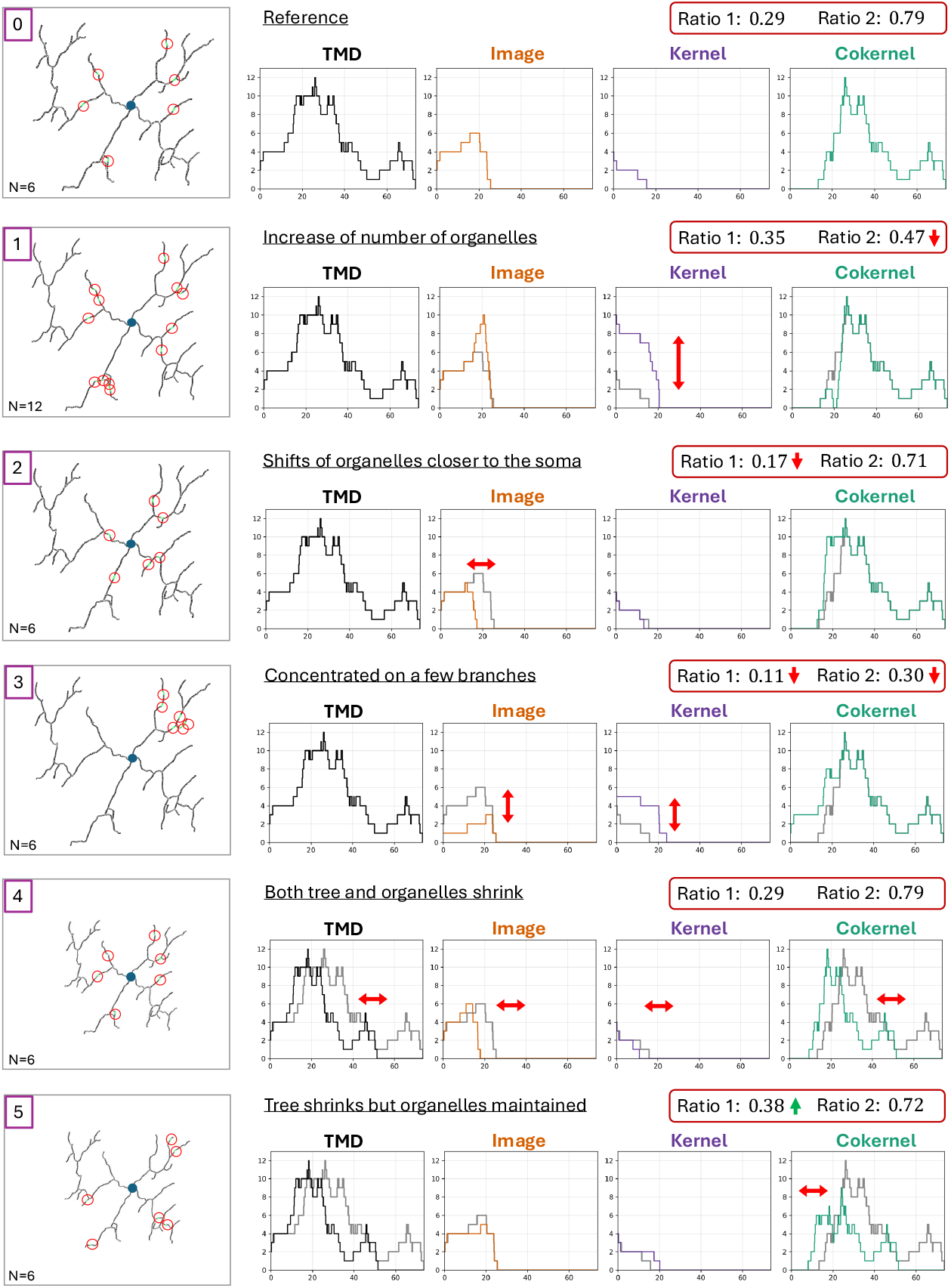
Representative examples of organelle distributions within a microglia tree. First column: microglia tree from **Figure 4** with synthetic distribution of organelles (red circles). In rows 0-3 the tree is maintained at its original scale; in rows 4-5 the scale is reduced by 30%. Number of organelles indicated in left bottom corner. Following columns: *Betti curves* of TMD (topological morphological descriptor, black) and the three invariants of the *chromatic TMD* : *image* (orange), recording the presence of organelles on branches, *kernel* (purple), capturing multiple organelle domains within individual branches, and *cokernel* (green), representing portions of the arbor that remain organelle-free, and *chromatic TMD ratios*. The first row serves as a reference for the following representative examples. Reference Betti curves indicated in grey. Notable changes in Betti curves and ratios indicated with red arrows.

**Figure S2:**
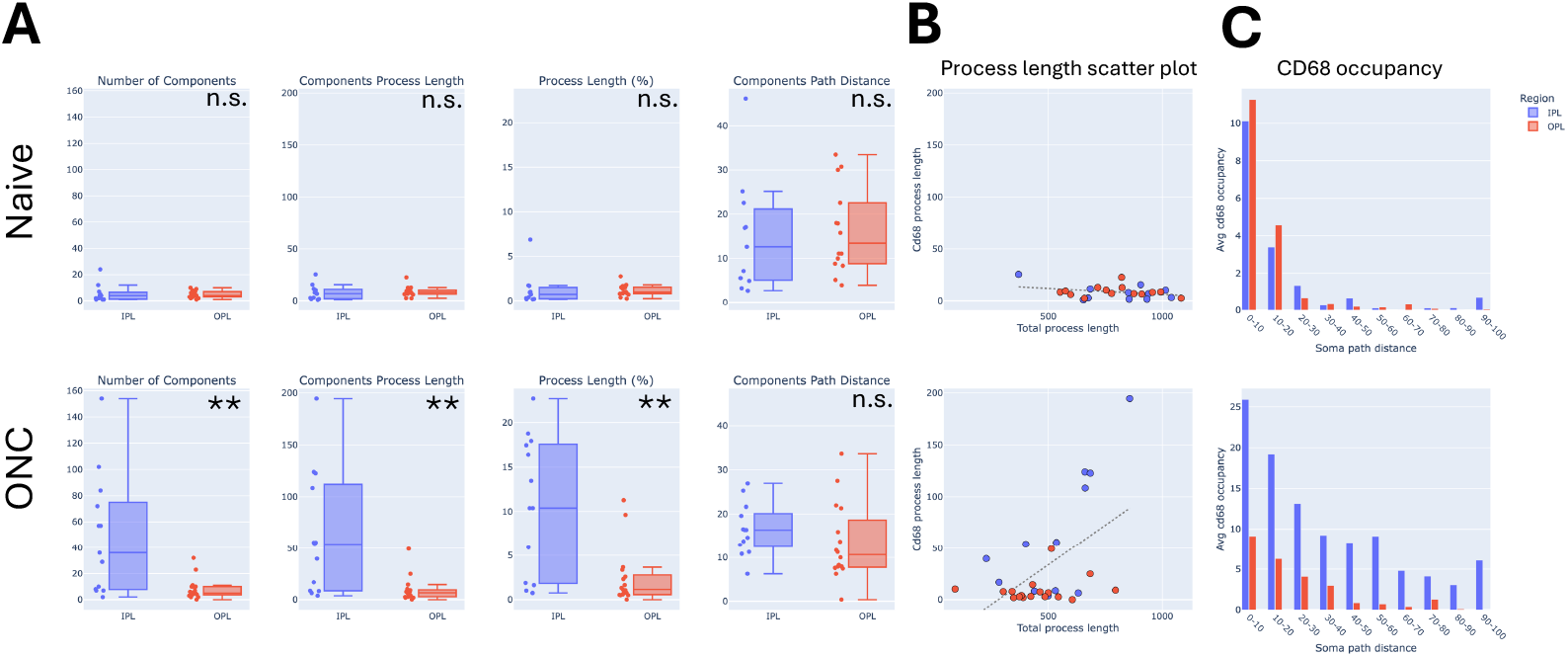
Morphometric and occupancy-based analyses support CD68 reorganization in the IPL. **A**. Box plots of CD68 *number of components, components process length, process length in %* and *components path distance* for naive (top) and optic nerve crush (ONC, bottom) with central line as median with interquartile range (Q1–Q3), with whiskers extending to the most extreme non-outlier values (1.5×IQR). All individual observations are overlaid, t-tests * *p <* 0.05, ** *p <* 0.01. **B**. Scatter plot of total process length *versus* process length covered by CD68, each dot represents a single microglia, for naive (top) and ONC (bottom) conditions, color-coded by region (blue for OPL and red for IPL). Dashed regression line. **C**. CD68 occupancy as percentage of process length at binned path distances to the soma in naive (top) and ONC (bottom) conditions, color-coded by region.

**Figure S3:**
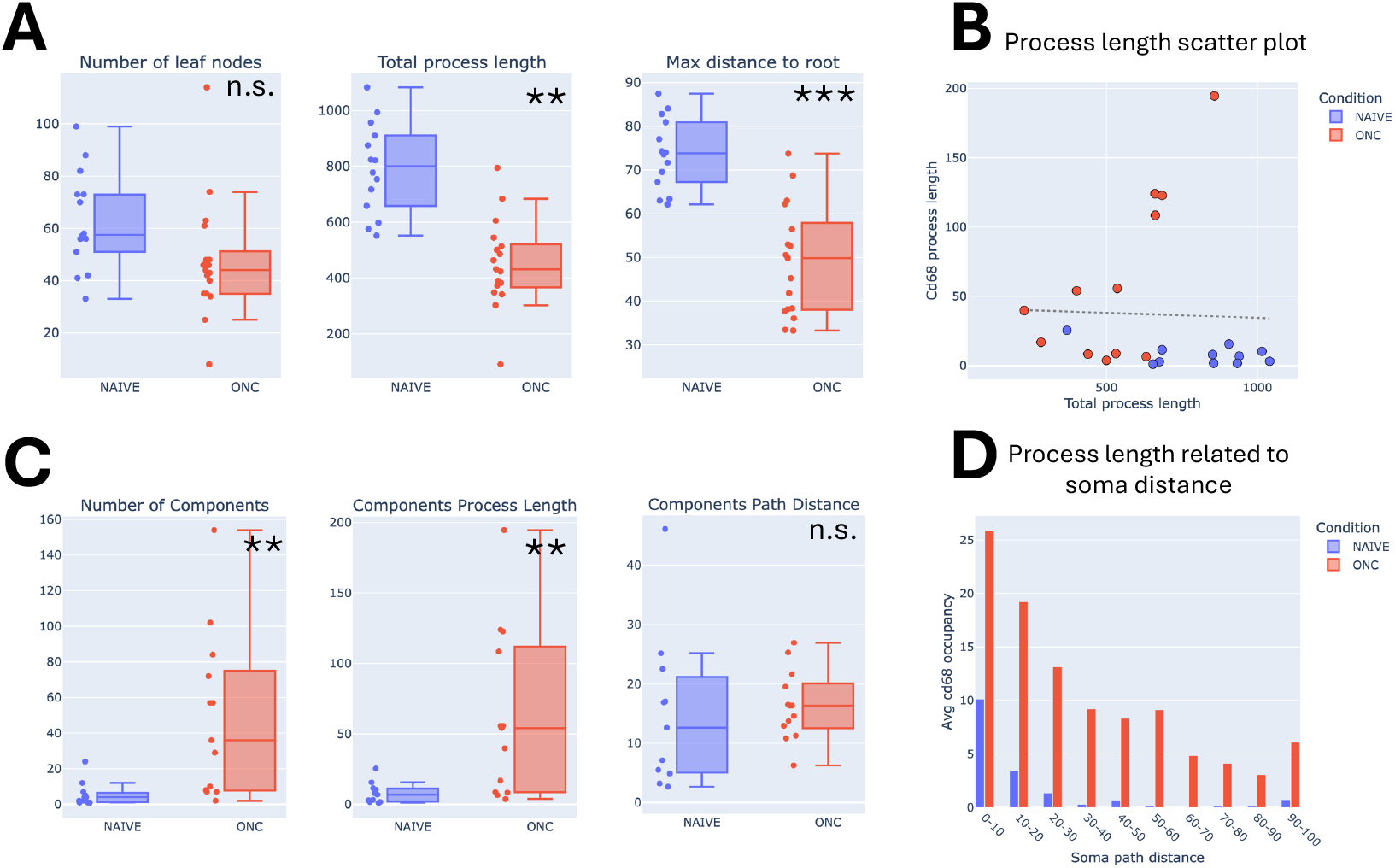
Morphometric and occupancy-based analyses support morphological adaptation with preserved CD68 constraints in IPL. **A**. Box plots of *number of leaf nodes, total process length* and *maximum distance to root (soma)* with central line as median with interquartile range (Q1–Q3), with whiskers extending to the most extreme non-outlier values (1.5×IQR). All individual observations are overlaid, t-tests ** *p <* 0.01, *** *p <* 0.001. **B**. Scatter plot of total process length *versus* process length covered by CD68, each dot represents a single microglia, color-coded by condition (blue for naive and red for ONC). Dashed regression line. **C**. Box plots of CD68 *number of components, components process length, process length in %* and *components path distance* with central line as median with interquartile range (Q1–Q3), with whiskers extending to the most extreme non-outlier values (1.5×IQR). All individual observations are overlaid, t-tests * *p <* 0.05, ** *p <* 0.01, *** *p <* 0.001. **D**. CD68 occupancy as percentage of process length at binned path distances to the soma, color-coded by condition.

**Figure S4:**
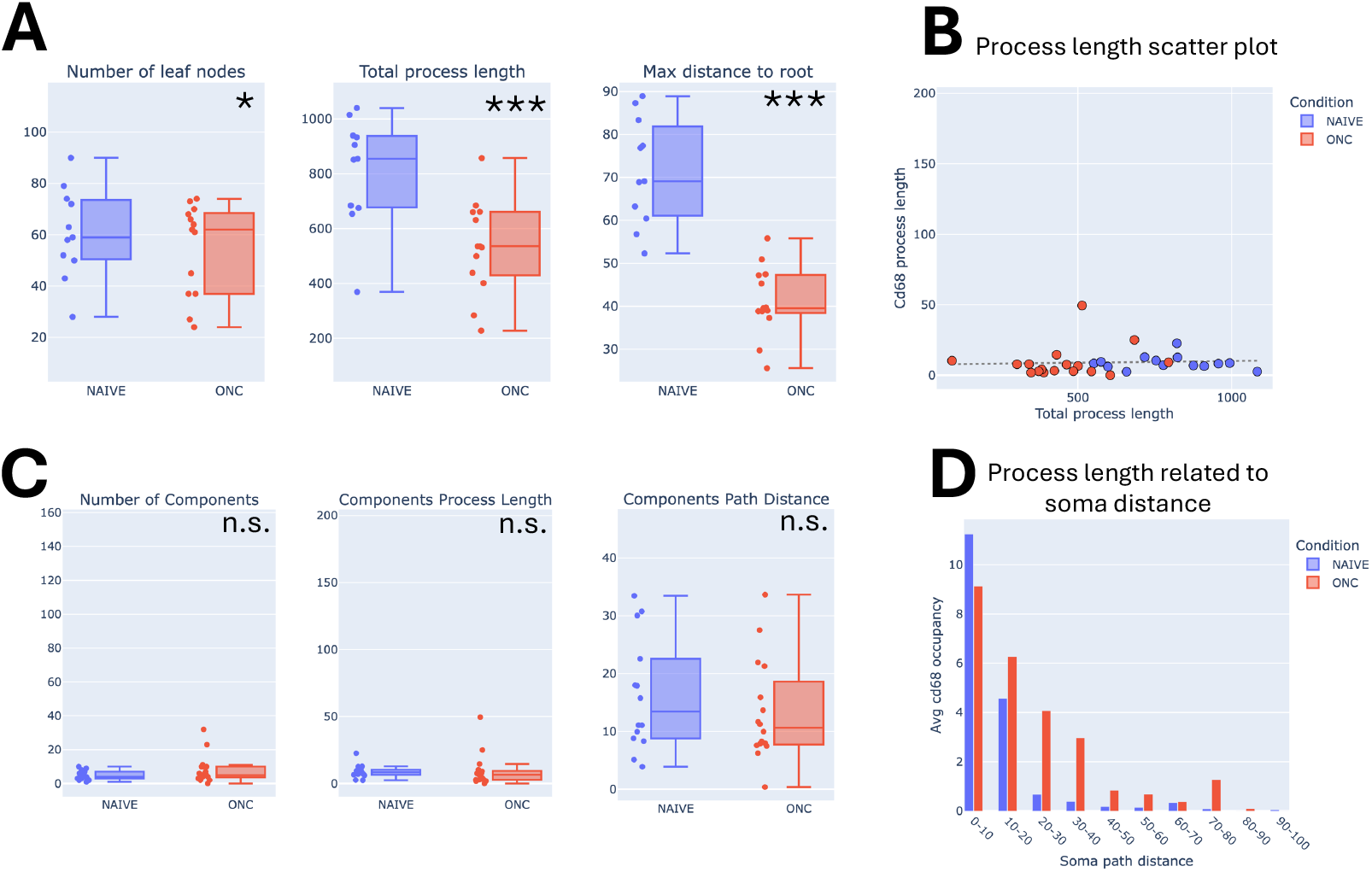
Morphometric and occupancy-based analyses support morphological adaptation with preserved CD68 constraints in OPL. **A**. Box plots of *number of leaf nodes, total process length* and *maximum distance to root (soma)* with central line as median with interquartile range (Q1–Q3), with whiskers extending to the most extreme non-outlier values (1.5×IQR). All individual observations are overlaid, t-tests * *p <* 0.05, *** *p <* 0.001. **B**. Scatter plot of total process length *versus* process length covered by CD68, each dot represents a single microglia, color-coded by condition (blue for naive and red for ONC). Dashed regression line. **C**. Box plots of CD68 *number of components, components process length, process length in %* and *components path distance* with central line as median with interquartile range (Q1–Q3), with whiskers extending to the most extreme non-outlier values (1.5×IQR). All individual observations are overlaid, t-tests * *p <* 0.05, ** *p <* 0.01, *** *p <* 0.001. **D**. CD68 occupancy as percentage of process length at binned path distances to the soma, color-coded by condition.

**Figure S5:**
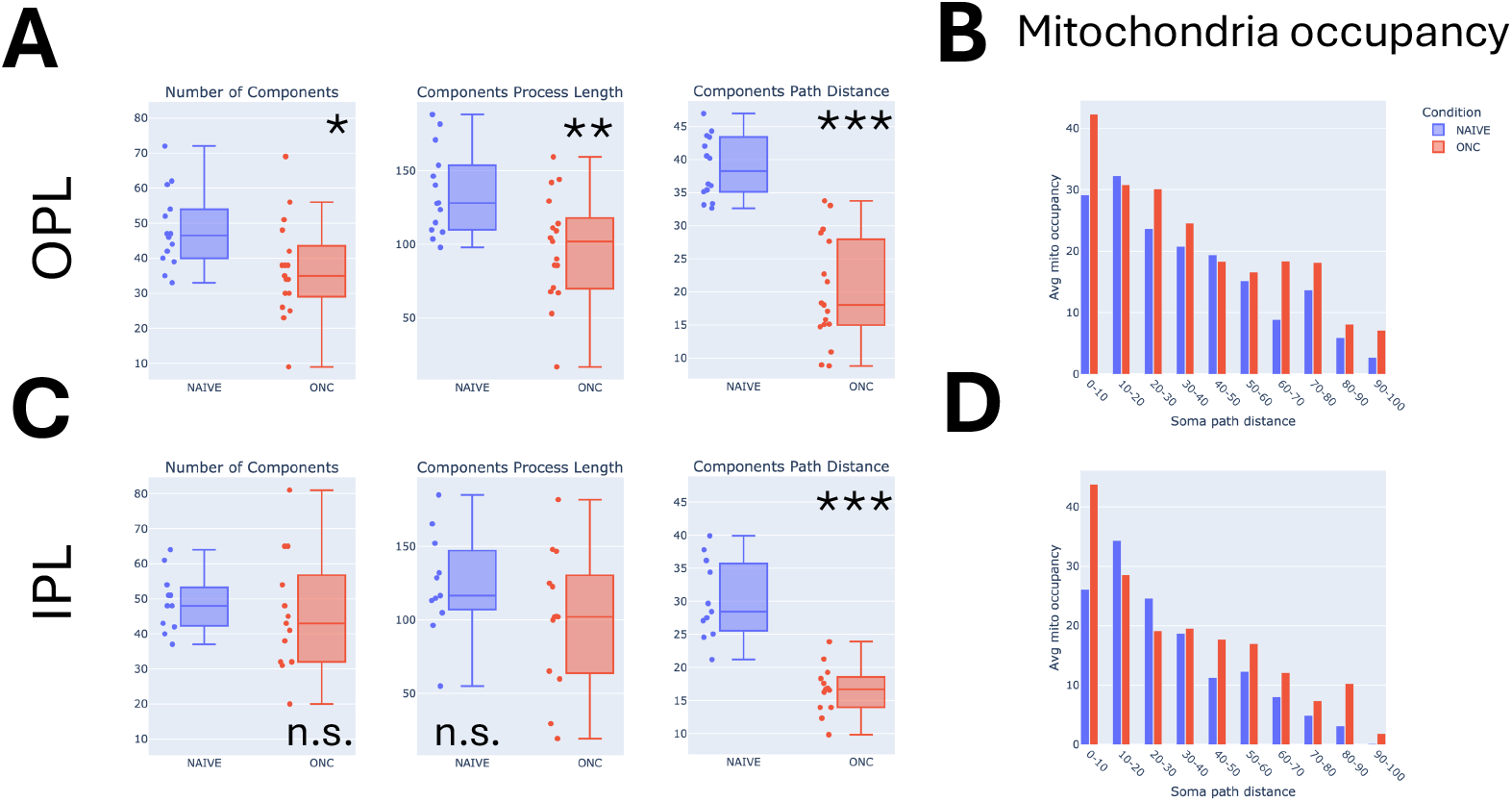
Morphometric and occupancy-based analyses for mitochondria. **A**,**C**. Box plots of *number of components, components process length* and *components path distance* for OPL (**A**) and IPL (**C**) with central line as median, interquartile range (Q1–Q3), and whiskers extending to the most extreme non-outlier values (1.5×IQR). Dots next to boxplot, individual observations. t-tests, *** *p <* 0.001, ** *p <* 0.01, * *p <* 0.05, ^ns^ *p >* 0.05. **B**,**D**. Mitochondria occupancy as average percentage of process length at binned path distances to the soma in OPL (**B**) and IPL (**D**), color-coded to the condition.

**Figure S6:**
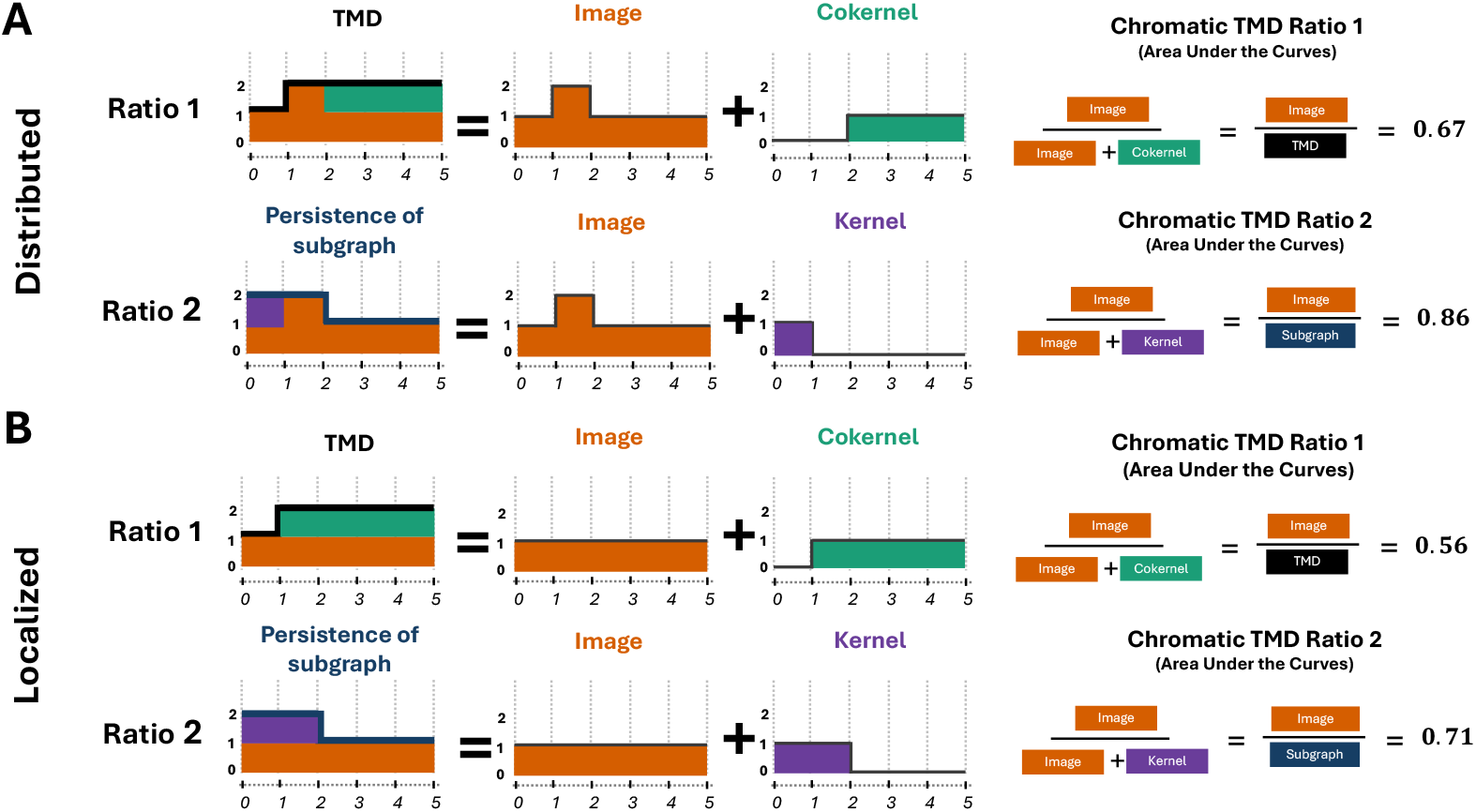
**A-B**. Derivation of *Chromatic TMD ratios* for the distributed (**A**) and localized (**B**) schematic trees from **Figure 3 A**. For *ratio 1* (**A** first row, **B** first row) the area under the TMD Betti curve (first column) is decomposed into the area under the *Image* Betti curve (second column, orange) and the area under the *cokernel* Betti curve (third column, green). Similarly, for *ratio 2* (**A** second row, **B** second row), the Betti curve for the persistence of subgraph is decomposed into the Betti curves of *image* (orange) and *kernel* (purple). The *distributed* tree had a higher *ratio 1* than the *localized* tree suggesting that more branches contain organelles. Also, *ratio 2* was higher in the *distributed* tree emphasizing that the organelles spread across the branches.

## Supplementary material

### A Dataset description - Sample and animal count

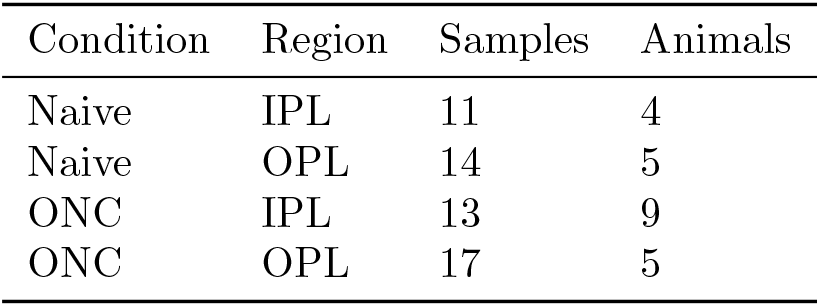

### B Algorithm

#### B.1 Setting

The algorithm takes as input the rooted tree *X* = (*V*_*X*_, *E*_*X*_), its subgraph *Y* = (*V*_*Y*_, *E*_*Y*_) and the function *f* : *X* → ℝ. It is initialized by being called on the root vertex.

For *w* ∈ *V*_*X*_ we call *X*(*w*) the subtree of *X* rooted at *w*. That is the tree containing *w* and all *v* s.t. *w < v*, as well as edges between these vertices. And *Y* (*w*) = *X*(*w*) ∩ *Y* .

We consider a vertex *w* with *n* children *v*_1_, …, *v*_*n*_. If *n* = 0, *w* has no children and this constitutes the base case of the algorithm. In the recursive step, *n >* 0, the input is *X*(*w*), *Y* (*w*) and the restriction of *f* to *X*(*w*), denoted by *f* |_*X*(*w*)_.

We let *s* = *f* (*w*). If *n >* 0 we let *t* := min {*f* (*v*_1_), …, *f* (*v*_*n*_))} and if *n* = 0 we can choose any *t > s*. We are interested in the following diagram:

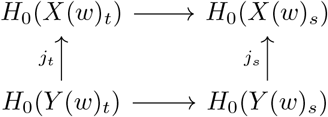

Note that *X*(*w*)_*s*_ = *X*(*w*) and 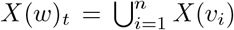. Since *X*(*v*_*i*_), *X*(*v*_*j*_) are disjoint for all *i, j* = 1, …, *n, i /*= *j* we have 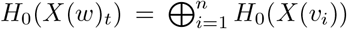 and 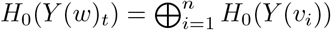. The persistence barcodes for *image, kernel, cokernel* corresponding to *X*(*w*), *Y* (*w*) can thus be fully described in terms of the barcodes *image, kernel, cokernel* of *X*(*v*_*i*_), *Y* (*v*_*i*_) for *i* = 1, …, *n* together with the transition *t* ≤ *s* described in the recursive step.

Note also that if *w* is the root *X*(*w*)_*t*_ = *X* and *Y* (*w*)_*t*_ = *Y*, thus we recover the barcode decomposition for *X, Y* by calling the algorithm on the root vertex.

#### B.2 Base case

In the base case the algorithm is called on a vertex *w* with no children (*n* = 0). If *w*∉ *V*_*Y*_ the diagram takes the following form.

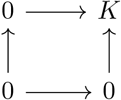

From this we see that *image*_*t*≤*s*_ and *kernel*_*t*≤*s*_ are the zero map, while *cokernel*_*t*≤*s*_ is 0 → *K*. Thus a bar starts at *s* in the *cokernel*.

If *w* ∈ *V*_*Y*_ the diagram takes the following form.

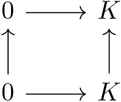

From this we see that *image*_*t*≤*s*_ is 0 → *K* while *kernel*_*t*≤*s*_ and *cokernel*_*t*≤*s*_ are the zero map. Thus a bar starts at *s* in the *image*.

#### B.3 Recursion step

We now consider a vertex *w* with *n* children *v*_1_, …, *v*_*n*_. The following variables will be used in the algorithm:

- 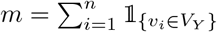, that is, the number of children of *w* that are in *V*_*Y*_ .
- 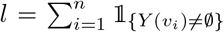, that is the number of children *v* of *w* such that at least one vertex of the subtree *X*(*v*) belongs to *V*_*Y*_ .

We split the exposition into four cases, depending on whether *ℓ* = 0 or *ℓ >* 0 and whether *w*∉ *V*_*Y*_ or *w* ∈ *V*_*Y*_ .

**Figure S7:**
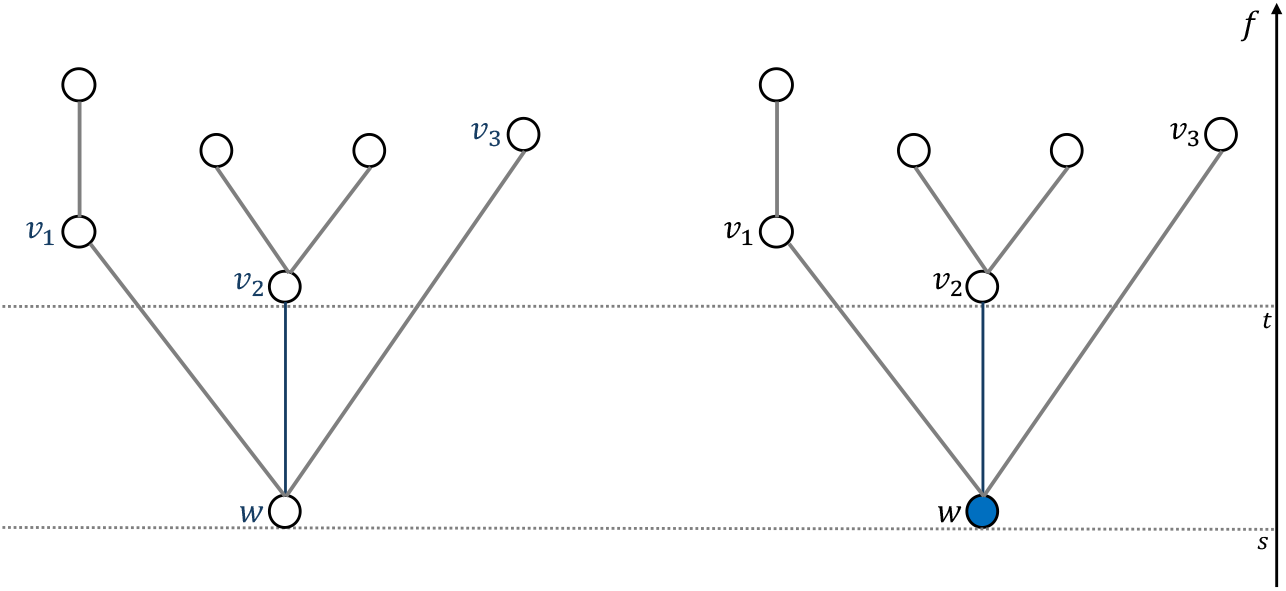
Illustration of Cases 1 and 2 (*ℓ* = 0). Transition from *t* = min*{f* (*v*_1_), …, *f* (*v*_*n*_)*}* to *s* = *f* (*w*). Vertices and edges belonging to *Y* are indicated in blue. Here *ℓ* = 0 and thus none of the subtrees rooted at *v*_1_, *v*_2_, *v*_3_ have any vertices belonging to *V*_*Y*_ . **Left:** *w*∉ *V*_*Y*_ . **Right:** *w* ∈ *V*_*Y*_ .

##### B.3.1 Case 1: *ℓ* = 0, *w*∉ *V*_*Y*_

This case is illustrated in Figure S7 and the commutative diagram takes the following form.

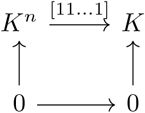

From the diagram we see that *image*_*t*≤*s*_ and *kernel*_*t*≤*s*_ are 0 → 0, while *cokernel*_*t*≤*s*_ 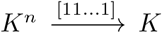 *K*, or by a change of basis 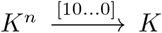. Thus by the elder rule convention we let all bars but the bar with the largest birth time stop at time *s* (if there are several such bars with the same birth time, one is chosen at random).

##### B.3.2 Case 2: *ℓ* = 0, *w* ∈ *V*_*Y*_

This case is illustrated in Figure S7 and the commutative diagram takes the following form.

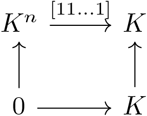

From the diagram we can see that *image*_*t*≤*s*_ is 0 → *K*, thus a bar starts at *s* in the *image, kernel*_*t*≤*s*_ is 0 → 0 and *cokernel*_*t*≤*s*_ is *K*^*n*^ → 0 thus all bars in the *cokernel* stops at time *s*.

##### B.3.3 Case 3: *ℓ >* 0, *w*∉ *V*_*Y*_

**Figure S8:**
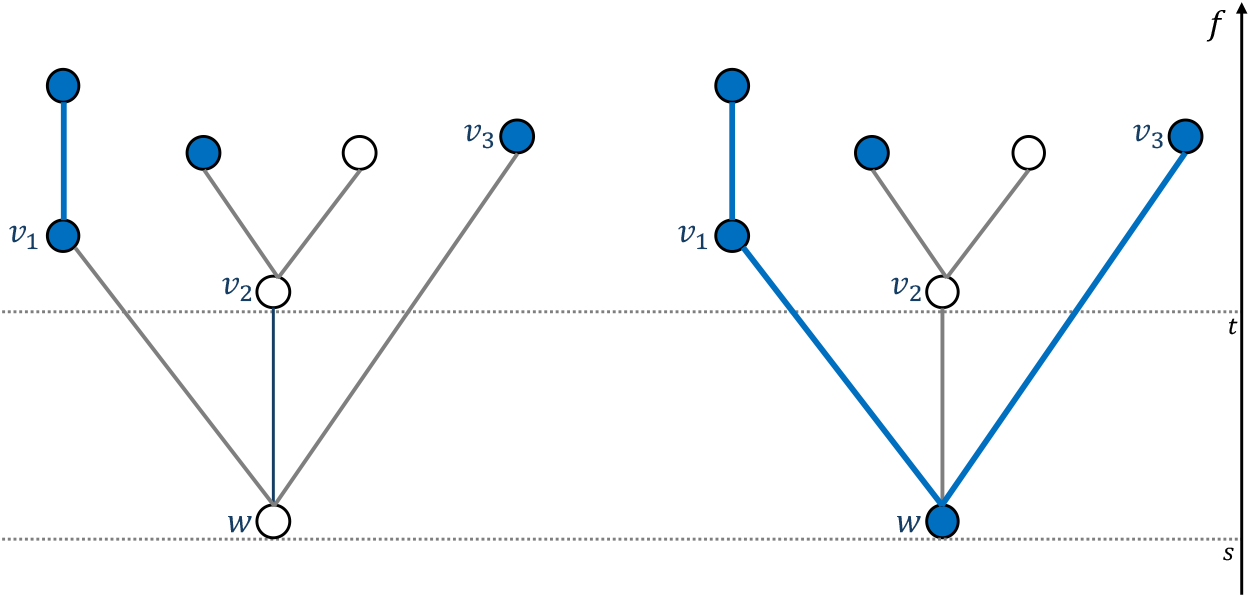
Illustration of Cases 3 and 4 (*ℓ >* 0). Transition from *t* = min*{f* (*v*_1_), …, *f* (*v*_*n*_)*}* to *s* = *f* (*w*). Vertices and edges belonging to *Y* are indicated in blue. Here *ℓ* = 3 (all the subtrees rooted at *v*_1_, *v*_2_, *v*_3_ have nodes belonging to *V*_*Y*_) and *m* = 2 (since *v*_1_, *v*_3_ ∈ *V*_*Y*_). **Left:** *w*∉ *V*_*Y*_ . **Right:** *w* ∈ *V*_*Y*_ .

This case is illustrated in Figure S8 and the commutative diagram takes the following form.

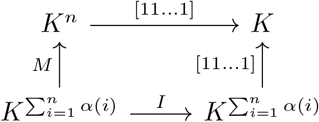

In the diagram *α*(*i*) denotes the number of connected components of *Y* (*v*_*i*_) for *i* = 1, …, *n, I* is the identity matrix and *M* is as below:

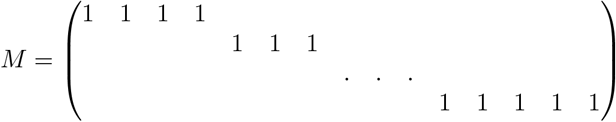

That is, on the first row *M* has 1 in the columns 1, …, *α*(1) and 0 elsewhere. In rows 2, …, *n, M* has 1 in column 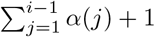 to column 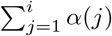.

From the diagram we see that *image*_*t≤s*_ is 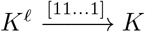, or by a change of basis 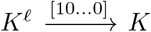. Thus by the elder rule convention we let all bars but the bar with the largest birth time stop at time *s* (if there are several such bars with the same birth time, one is chosen at random).

The *kernel* of *j*_*t*_ is thus of dimension 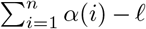. While the *kernel* of *j*_*s*_ is of dimension 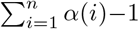 By commutativity of the diagram we have *kernel*(*j*_*t*_) ⊂ *kernel*(*j*_*s*_). Thus we have that *ℓ* − 1 bars start in the *kernel* at time *s*.

Finally *cokernel*_*t*≤*s*_ is *K*^*n*−*ℓ*^ → 0 thus all bars in the *cokernel* stop at time *s*.

##### B.3.4 Case 4: *ℓ >* 0, *w* ∈ *V*_*Y*_

This case is illustrated in Figure S8 and the commutative diagram takes the following form.

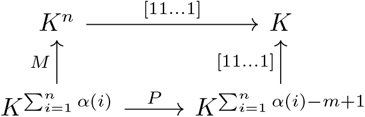

The matrix *P* now depends on the variable *m*. If *m* = 1 *P* is the identity matrix as in Case 3.

If *m* = 0 we have that *P* is the identity matrix with a row with all zeros added:

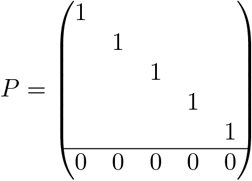

If *m >* 1 we can write *P* on the form:

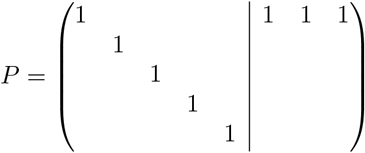

That is, for the first 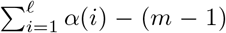 columns we have the identity map and the next *m* − 1 columns we have 1 on the first row (corresponding to the components in *Y* (*w*) being merged).

From this we see that *image*_*t≤s*_ is 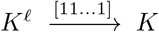, or by a change of basis 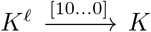. Thus by the elder rule convention we let all bars but the bar with the largest birth time stop at time *s* (if there are several such bars with the same birth time, one is chosen at random).

The *kernel* of *j*_*t*_ is thus of dimension 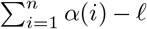. While the *kernel* of *j*_*s*_ is of dimension 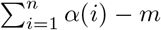. By commutativity of the diagram we have that the image of *kernel*(*j*_*t*_) by *P* must be contained in *kernel*(*j*_*s*_). Thus we have that *ℓ* − *m* bars start in the *kernel* at time *s*. Finally *cokernel*_*t*≤*s*_ is *K*^*n*−*ℓ*^ → 0 thus all bars in the *cokernel* stop at time *s*.

###### Algorithm 2

ChromaticTMD

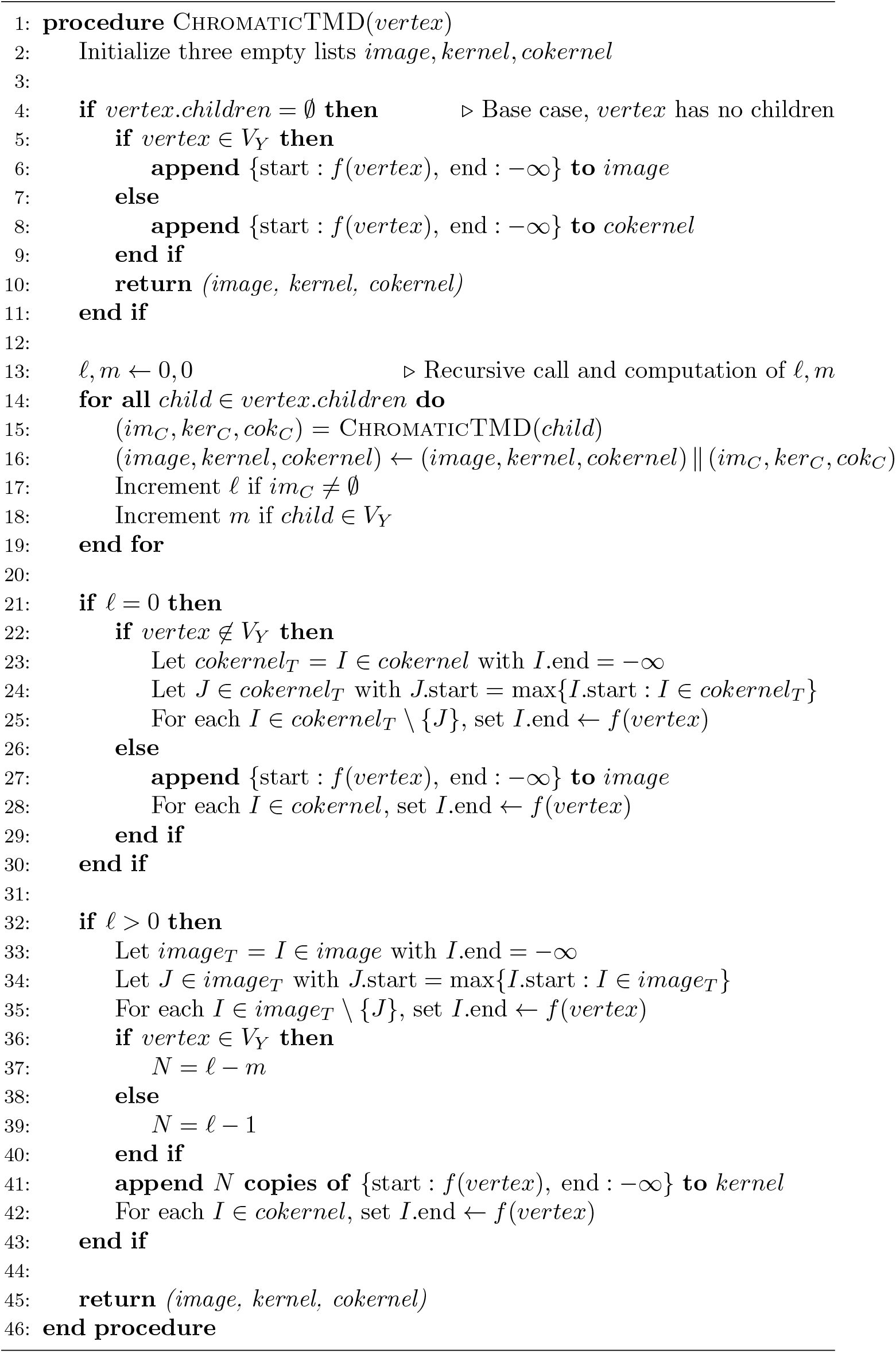

